# Systematic profiling of the acetyl lysine machinery reveals a role for MAPKAPK2 in bromodomain inhibitor resistance

**DOI:** 10.1101/2024.07.22.604604

**Authors:** Pata-Eting Kougnassoukou Tchara, Jérémy Loehr, Lucas Germain, Zachary Steinhart, Beatriz Gonzalez-Badillo, Anahita Lashgari, François J.M. Chartier, Monika Tucholska, Sarah Picaud, James D.R. Knight, Stéphane Angers, Nicolas Bisson, Colin R. Goding, Étienne Audet-Walsh, Panagis Filippakopoulos, Anne-Claude Gingras, Jean-Philippe Lambert

## Abstract

Bromodomain (BRD)-containing proteins are chemically tractable multi-domain scaffolding molecules involved in acetyl lysine (Kac) signaling. BRD inhibitors have shown promise in clinical oncology, including melanomas; however, their narrow therapeutic windows and issues with resistance in pre-clinical models highlight the need to better understand the functions of and interconnection between BRD-containing proteins. Here, we use complementary interaction-mapping techniques (affinity purification and proximity-dependent biotinylation) to map the interactions of 39 of the 42 BRD-containing proteins and 110 additional proteins that physically or functionally associate with them. We uncover 3,892 novel interactions and reveal the intricate connectivity of the Kac machinery. Chemical inhibition of multiple BRD classes revealed that inhibiting BETs—but not mSWI/SNF or CREBBP/EP300 proteins—dramatically rewired the interactome. Finally, we identified MAPKAPK2 activity as a critical determinant of BET inhibitor sensitivity in melanoma through its impact on chromatin composition remodeling.

**In Brief:** Kougnassoukou Tchara *et al*. generate a static protein interaction map of the human acetyl lysine machinery by coupling two complementary functional proteomics approaches (FLAG affinity purification and proximity-dependent biotinylation) to mass spectrometry. They also investigate network changes upon bromodomain inhibition, and describe a novel resistance mechanism mediated by the p38 stress signaling pathway that causes significant metabolic changes.

**Highlights:**

- Two complementary interaction proteomics analyses of the human acetyl lysine machinery were performed.
- Novel target- and compound-specific impacts of bromodomain inhibitors were identified.
- *MAPKAPK2* was identified as a novel resistance gene to BET bromodomain inhibitors in melanoma.
- BET bromodomain inhibition leads to metabolic adaptation in melanoma.

## Introduction

Lysine acetylation (Kac) is a key post-translational modification regulating chromatin function (Kouzarides, 2000), in addition to numerous facets of metabolism and cell signaling (Verdin and Ott, 2015). Kac neutralizes the positive charge of the lysine side-chain with several mechanistic consequences, including chromatin structure (Grunstein, 1997), protein binding (Yuan et al., 2005), catalytic activity (Sabo et al., 2008), and the deposition of other lysine modifications (Min et al., 2010). Importantly, Kac can also mediate specific protein-protein interactions upon recognition by conserved protein domains called bromodomains (BRDs), which primarily bind Kac residues on histones (Dhalluin et al., 1999; Jacobson et al., 2000; Owen et al., 2000). Humans encode 61 distinct BRDs in 42 proteins. We previously reported the structural determinants leading to BRD-mediated Kac recognition on histones, as well as over 1,000 novel sites (mostly on histones), including multiple adjacent sites recognized by single BRDs (Kougnassoukou Tchara et al., 2019; Lambert et al., 2019; Owen et al., 2000). However, as these studies depended on isolated domains outside their cellular contexts, they identified only small fractions of the thousands of reported Kac sites (Choudhary et al., 2009; Kim et al., 2006), providing a partial view of BRD specificity. Interest in improving our understanding of BRD functions and specificity has greatly increased following the discovery that BRD-Kac interactions are druggable (Filippakopoulos et al., 2010; Nicodeme et al., 2010), making them attractive for pharmaceutic intervention. Multiple compounds with varied specificities toward BRDs are now available, and several have shown promise in preclinical and phase I/II clinical studies on cancer and other health conditions (Ameratunga et al., 2020; Cescon et al., 2024; Fujisawa and Filippakopoulos, 2017; Hamilton et al., 2023).

Importantly, BRD-containing proteins contain additional modular protein interaction domains that can help recruit other proteins to the acetylated protein target, enabling them to contribute to complex processes such as chromatin remodelling and transcription. While many interactions have been reported for individual BRD-containing proteins using different systems and experimental approaches, a systematic survey using standardized reagents remains to be performed. This prevents assessment of the relative specificities and biological effects of small molecule BRD inhibitors—a major gap in our understanding of the specificity and scaffolding functions of the acetylation recognition machinery. Here, we investigate the functional interactions of 39 BRD family members and 100 BRD-associated proteins to reveal a network of > 2,000 interacting proteins. This rich interactome allowed us to identify MAPKAPK2 as a significant kinase contributing to bromo-and-extra-terminal (BET)-family BRD inhibitors sensitivity in melanoma.

## Results

### The interaction landscape of human bromodomain-containing proteins

To better understand how protein-protein interactions contribute to Kac signaling, we employed proteomics approaches to systematically identify the interactors of each human BRD-containing protein. We first assembled a near-complete collection of full-length constructs for 39 of the 42 human BRD-containing proteins (**Table S1**) as 3×FLAG and BirA*-FLAG expression vectors for affinity purification coupled to mass spectrometry (AP-MS) and proximity-dependent biotinylation (BioID)(Roux et al., 2012), respectively. Each vector was stably transfected in the isogenic tetracycline-inducible Flp-In T-REx HEK293 system (Roux et al., 2012). HEK293 cells express mRNAs encoding most BRD-containing proteins (except BRDT, TAF1L, and TRIM66), acetyltransferases and deacetylases, making them suitable for our analyses (**Figure S1A-B**).

AP-MS and BioID are orthogonal approaches that, when combined, provide a broader view of the interactome (Lambert et al., 2015). In AP-MS, the bait of interest is fused to an epitope tag (3×FLAG here), enabling purification of the bait and its interactors on an anti-tag resin. Here, we used AP-MS conditions optimized for proteins associated with the chromatin (Lambert et al., 2014), where most of the Kac machinery is found (Gong et al., 2015). In BioID, the bait is expressed as a fusion with an abortive biotin ligase (BirA*) that activates exogenous biotin (added to the cell medium) in its vicinity (Roux et al., 2012). Proteins proximal to the bait in its cellular context become covalently biotinylated, enabling their subsequent recovery using streptavidin reagents. Since the proximal interactions do not need to be maintained following cell lysis, BioID is compatible with harsh lysis and purification conditions. As with AP-MS, we have extensively optimized the BioID protocol for use with chromatin-associated proteins (Lambert et al., 2019; Lambert et al., 2015).

To ensure data robustness, we analyzed two biological replicates for each bait by both AP-MS and BioID alongside negative controls (see Supplemental Experimental Procedures). Potential interactions were scored using Significance Analysis of INTeractome (SAINT)express (Teo et al., 2014), averaging the SAINT scores across replicates before Bayesian false discovery rate (FDR) estimation to ensure that only interactions detected in both replicates were reported. Interactions passing the estimated 1% FDR threshold were deemed high-confidence, and scored interactions were quantitatively reproducible across biological replicates (*R^2^* of 0.8–0.84, **Figure S1C, D**). We identified 668 and 1,046 high-confidence interactions for the 39 BRD-containing proteins using AP-MS and BioID, respectively (**Figure S1E**). Individually, BRD-containing proteins, from all BRD families we had previously described (Filippakopoulos et al., 2012), displayed 0–100 high-confidence AP-MS and 0–103 high-confidence BioID proximal interactions, with medians of 7 and 20, respectively (**Figure 1A**; **Table S2A, S2B**).

**Figure 1.**
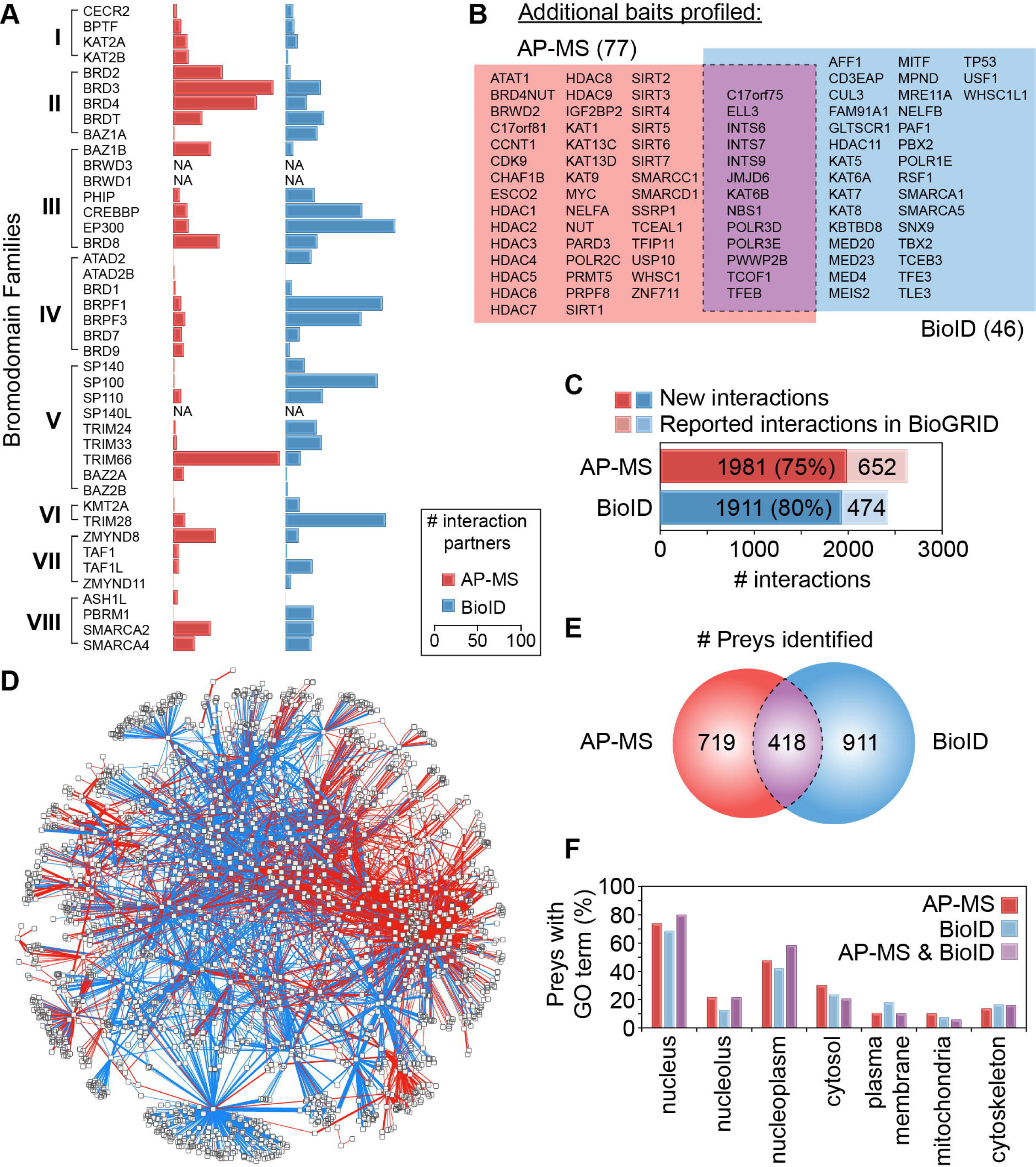
– A static interactome of human bromodomain-containing and -associated proteins. (**A**) Interaction partners identified for the BRD-containing proteins ordered by BRD family (as defined in (Filippakopoulos et al., 2012)) in Flp-In T-REx HEK293 cells by AP-MS (red) and BioID (blue). (**B**) Additional baits profiled by AP-MS (red), BioID (blue), or both approaches (purple). (**C**) Comparison of protein-protein or proximal interactions detected here with those reported in BioGRID. (**D**) An edge-weighted force-directed layout of all significant (SAINTexpress ≤ 1% FDR) AP-MS (red edges) and BioID (blue edges) interactions identified in the systematic screen. Edge thicknesses represent the average numbers of spectral counts detected for each prey. (**E**) Overlap between the preys detected by AP-MS (red) and BioID (blue). (**F**) Proportions of preys identified by AP-MS (red), BioID (blue), or both approaches (purple) associated with the indicated cellular compartment GO terms.

To help validate the interactome and better understand its modularity, we selected 37 high-confidence interactors (or proximal interactors) of BRD-containing proteins for reciprocal interaction proteomics (of these additional baits, 15 (40.5%) recovered the initial BRD-containing proteins). We also included 53 baits functionally related to the Kac machinery, including most lysine acetyltransferases and deacetylases and many known transcriptional regulators (**Table S1A**). These efforts identified an additional 1,955 preys for 77 baits by AP-MS and 1,340 preys for 46 baits by BioID. Collectively, our analyses yielded 5,009 high-confidence interactions involving 2,057 distinct proteins (**Table S2A**, **S2B**), 3,892 of which are not previously reported in BioGRID (Oughtred et al., 2019) (∼78%; **Figure 1C**).

The AP-MS and BioID datasets revealed roughly the same proportions of previously reported interactions (**Figure 1C**), confirming that both provide valid views of the acetylation machinery interactome. However, despite their similar recall, the BioID and AP-MS results only modestly overlapped. An edge-weighted force-directed layout of the interaction network generated in Cytoscape (Shannon et al., 2003) revealed segregation of the AP-MS and BioID data (**Figure 1D**), and only 339 (19.7%) of the 1,721 preys identified with the 73 baits profiled with both approaches were shared between them (**Figure 1E**). This highlights the complementarity of these datasets, as we previously observed for histones (Lambert et al., 2015). Consistent with the reported localizations of most Kac machinery components (Kim et al., 2006), gene ontology (GO) cellular compartment term analysis confirmed that most high-confidence interaction partners localized to the nucleus (**Figure 1F**).

### The Kac machinery shows high interconnectivity

Globally, our Kac interactome was enriched for known acetylated proteins and acetylation sites (Hornbeck et al., 2019) compared to the HEK293 proteome and datasets acquired with the same approaches (*e.g*., (St-Denis et al., 2016)) (**Table S2C**), suggesting high connectivity in the acetylation machinery. Since each BioID bait reports on its proximal interactors, the correlation profiles between preys localized near baits can provide insights into the organization of the endogenous proteins (see **Experimental Procedures**; (Go et al., 2021; Gupta et al., 2015; Youn et al., 2018)). Pearson correlation analysis of the prey-prey profiles in the BioID data uncovered several clusters of correlated preys defined by proximal interactions with one or more Kac components (**Figure 2A**, **Table S2D**). For example, *cluster 1* contained Cullin 4 E3 ligase (CUL4) and COP9 signalosome components, which were connected primarily to the BRD-containing protein PH-interacting protein (PHIP); this bait was also a prey of several other baits in our network (**Figure 2B**). *Cluster 2* was characterized by associations made by the transcription intermediary factor 1 (TIF1) family of BRD proteins (TRIM24, TRIM28, and TRIM33). These BRD-containing proteins are known to hetero-oligomerize (Germain-Desprez et al., 2003; Peng et al., 2002), and some of their preys were previously reported (reviewed in (Hatakeyama, 2011)). However, they also formed unique interaction networks with transcriptional regulators (TRIM24), KRAB domain-containing zinc finger proteins (TRIM28), and co-activators (TRIM33; **Figure 2C**). *Cluster 3* contained the SAGA and TIP60 lysine acetyltransferase complexes, which also formed novel interactions with transcription factors and BRD-containing proteins (**Figure 2D**).

**Figure 2.**
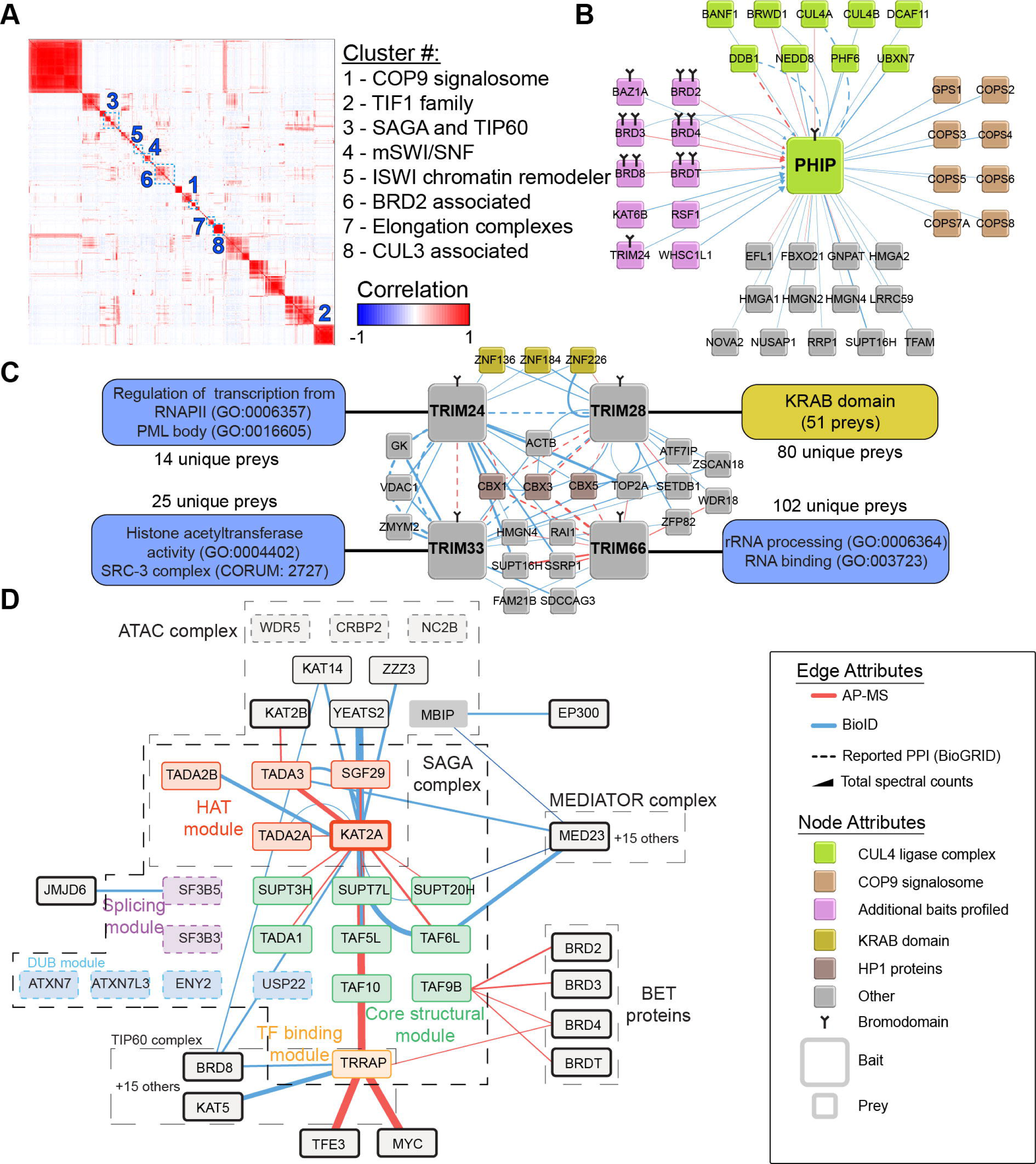
– High interconnectivity in the human Kac machinery. (**A**) Pearson correlations between prey protein profiles in the BioID dataset. See **Table S2D** for the cluster compositions. (**B–D**) Interaction networks for (**B**) PHIP, (**C**) members of the transcription intermediary factor 1 (TIF1) BRD family, and (**D**) the SAGA and TIP60 lysine acetyltransferase complexes. In **D**, undetected complex subunits are shown as transparent nodes.

Interconnectivity was also prevalent among the ATP-dependent chromatin remodeler mammalian (m)SWI/SNF complexes (**Figure 2A**, *cluster 4*), which associated with multiple BRD-containing proteins. AP-MS and BioID of each ATPase- and BRD-containing subunit (SMARCA2, SMARCA4, BRD7, BRD9, and PBRM1) recovered most of the mSWI/SNF subunits, as we recently reported (Agbo et al., 2023). Intriguingly, BRD9, a subunit of the ncBAF mSWI/SNF complex (Kadoch et al., 2013), was at the periphery of *cluster 4* which is consistent with its reported interactions with many other proteins, notably BRD4 and JMJD6 (Rahman et al., 2011).

*Cluster 5* contained all BRD-containing members of the ISWI chromatin remodeler family, including BAZ1A, BAZ1B, BAZ2A, BAZ2B, BPTF, and CECR2, and the two ATPase subunits (SMARCA1 and SMARCA5; **Figure S2A**). BAZ2B, the least characterized family member, recovered few interaction partners by AP-MS and BioID (**Figure S2A**). In systematic CRISPR/Cas9 screens in the Cancer Dependency Map (depmap.org; (Meyers et al., 2017)), BAZ2B was unique among ISWI chromatin remodelers in that its ablation promotes cancer cell proliferation (**Figure S2B**), suggesting a tumor suppressive role. This idea is supported by clinical cohorts, in which high BAZ2B expression was associated with better clinical outcomes for triple-negative breast cancer in The Cancer Genome Atlas’ pan-cancer analysis (**Figure S2C**). Therefore, we further investigated the BAZ2B interactome. As full-length BAZ2B has poor stability in HEK293 cells (*data not shown*), we expressed fragments containing only its BRD or PHD-BRD modules, fused to a nuclear localization sequence (NLS), which drastically enhanced its proximal interactions by BioID (**Figure S2D**). These included fundamental chromatin constituents (histones, high-mobility group proteins, *etc*.), transcriptional regulators (*e.g*., HDGF and PHF2), and proteins involved in DNA damage repair (*e.g*., TRIP12, RNF169). Adding an N2140F mutation to the BAZ2B fragments to prevent Kac anchoring within the BAZ2B BRD cavity dramatically reduced the levels of nuclear and chromatin-associated preys (**Figure S2D**). We also tested whether multiplexed constructs containing three copies of the BAZ2B BRD, which we previously showed to be effective H3K14ac masking agents (Savitsky et al., 2016), would enable effective interaction mapping, as recently reported (Villasenor et al., 2020). BAZ2B-3×BRD (with NLS) performed well in BioID experiments, further expanding its interactome. Mutating (N2140F) the BRDs abolished 30/63 interactions established by WT BAZ2B-3×BRD (SAINT FDR ≤ 1%) (**Table S2E**). Furthermore, all 68 novel interaction partners observed with BAZ2B-3×BRD^N2140F^-NLS were non-nuclear, demonstrating that a functional BRD is required for proper BAZ2B localization.

The last three clusters highlighted in **Figure 2A** were partly driven by their associations with BRD2, BRD4, and BETs (*clusters 6, 7, and 8*, respectively). *Cluster 6* was enriched for regulators of transcription initiation, including subunits of the transcription factor II D (TFIID) and mixed lineage leukemia complexes. *Cluster 7* contained most subunits of the little and super elongation complexes, which alleviate promoter-proximal RNAPII pausing (Smith et al., 2011a; Smith et al., 2011b). *Cluster 8* contained mainly cullin 3 (CUL3) adaptor proteins, which promote the ubiquitylation of various substrates (Pintard et al., 2004), and KBTBD8, whose associations with BRD2, BRD3, and BRD4 are drastically enhanced by BET BRD inhibitors (Lambert et al., 2019). Together, our interactome highlights the extensive involvement of BRD-containing proteins in many facets of the transcription cycle by RNAPII.

### BRD inhibitors have varying effects on Kac-associated interaction networks

The cellular impacts of small-molecule BRD inhibitors vary (Wu et al., 2019), and the precise molecular consequences of most remain undefined. We previously detailed how the pan-BET BRD inhibitor (+)-JQ1 (hereafter referred to as JQ1) drastically remodels the BET protein interaction networks (Lambert et al., 2019), raising the possibility that this is a general property of BRD inhibitors. To investigate this, we first focused on BRD inhibitors targeting the CREBBP/EP300 homologs, inhibitors known to modulate proliferation *in vitro* (Hay et al., 2014; Picaud et al., 2015). Without inhibition, AP-MS and BioID both recovered multiple known interactions for CREBBP and EP300, but with little overlap in the preys detected with each method (**Figure S3A-B**). In contrast to BET proteins (Lambert et al., 2019), the CREBBP and EP300 BioID interaction networks were mostly unaffected by I-CBP-112, SGC-CBP30, or the recently described KAT inhibitor A485 at dose preventing engagement of additional BRDs ((Lasko et al., 2017; Weinert et al., 2018); **Figure S3C; Table S2F**). Consistently, there were no major changes in the BioID interaction profiles of SMARCA2, SMARCA4, and PBRM1 after treatment with PFI3, a BRD inhibitor targeting mSWI/SNF BRDs ((Fedorov et al., 2015); **Figure S3D; Table S2G**). However, the BET BRD-inhibitor JQ1 modulated the proximal interactomes of SMARCA2 and SMARCA4 without engaging their BRDs (**Figure S3D**). These results suggest that BET BRDs are central nodes in the Kac machinery network, consistent with the dramatic interactome rewiring observed upon their inhibition.

To address this possibility directly, we systematically profiled the BRD4 AP-MS interaction network after treatment with six distinct BRD inhibitors that directly target either BRD4’s BRDs or those of one of its BRD-containing interactors (**Figure 3A-B; Table S2H**). Consistent with our previous report (Lambert et al., 2019), inhibitors directly targeting both BRD4 BRDs, such as JQ1 and the more potent ABBV-075 (Mivobresib; (McDaniel et al., 2017)), significantly reduced or enhanced its interactions (**Figure 3B**). ABBV-744, a BRD4 inhibitor that specifically binds its second BRD (Faivre et al., 2020), impacted BRD4’s interactome similarly to the pan-BRD BET inhibitor ABBV-075 tested (Pearson correlation 0.946), but with an overall smaller amplitude (**Figure 3A-3B**). Notably, only JQ1 induced neomorphic associations between BRD4, the centromeric protein CENPB, and the CENPA licensing complex MIS18 (MIS18A, MIS18B, and MIS18BP1), as recently reported by Corless *et al*. (Corless et al., 2023). Moreover, we identified a second putative JQ1-dependent BRD4-associated complex containing UACA, TRIOBP, and RAI14 (**Figure 3B**), which play roles in cytoskeleton dynamics. These results support the possibility that JQ1 stabilize and/or generate novel protein-protein interactions in addition to its BRD inhibition activity. Of note, this behavior appears to be unique to JQ1 among the BRD inhibitors that we tested. The CREBBP/EP300 BRD inhibitor SGC-CBP30 mildly impacted the BRD4 interactome, potentially due to its residual activity toward BET BRDs (Hay et al., 2014). Lastly, the BRD9 BRD inhibitor Bi-9564 and mSWI/SNF BRD inhibitor PFI3 did not significantly impact the BRD4 interactome (Pearson correlation versus DMSO of 0.931 and 0.887, respectively; **Figure 3B**). As most modulations of the Kac machinery network were generated by BET BRD inhibitors, we reasoned that refocusing on them would provide the best opportunity to identify co-targets that could be leveraged to improve BET BRD inhibitors’ clinical efficacy.

**Figure 3.**
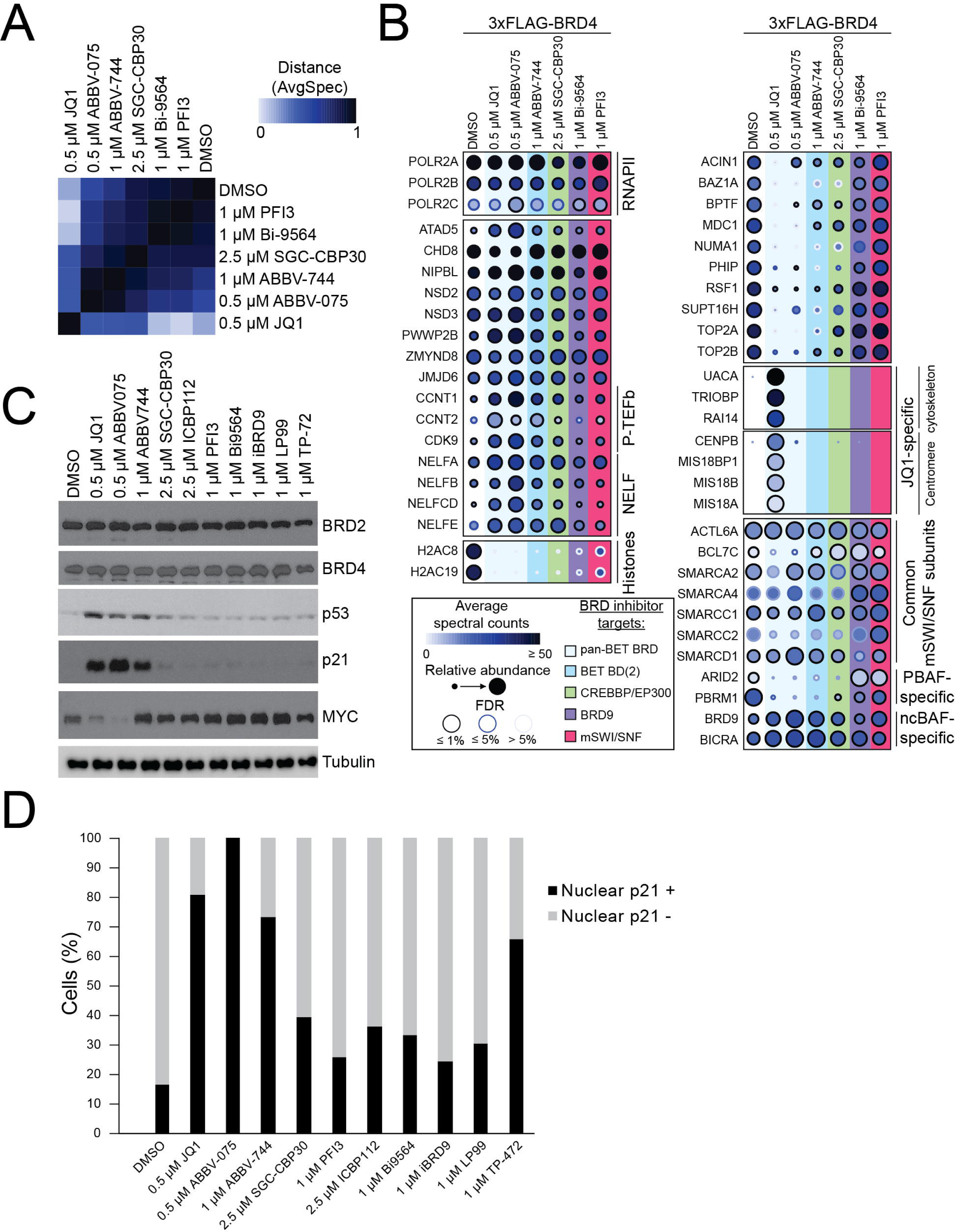
– Target- and compound-specific modulation of the BRD4 interaction network by BRD inhibitors. (**A**) Bait-bait Pearson correlations between BRD4 AP-MS samples from Flp-In T-REx HEK293 cells treated with the indicated BRD inhibitors for 1 h. (**B**) Dot plots of selected BRD4 interaction partners in Flp-In T-REx HEK293 cells. (**C**) Western blot analyses of the indicated proteins following 1-h treatments with the indicated BRD inhibitors in A375 cells. Tubulin was used as a loading control. (**D**) Quantification of cells treated with the indicated BRD inhibitors for 48 h displaying nuclear p21 using immunofluorescence. See **Figure S4A** for representative images.

In A375 melanoma cells, which are highly sensitive to BET inhibition (Zhao et al., 2018), BET inhibitor-induced changes in the BRD4 interactome were correlated with MYC downregulation and the upregulation of the tumor suppressor TP53 (p53) and its downstream effector CDKN1A (p21) (**Figure 3C**). When melanoma cells enter a more differentiated state, they display extended morphologies with long protrusions (Hoek and Goding, 2010). BET inhibitors induced this phenotype in A375 cells (**Figure S4A**), as initially reported by Segura *et al*. (Segura et al., 2013); however, the other BRD inhibitors tested did not (**Figure S4A**). As the master transcriptional regulator of melanocytes, MITF, impacts melanoma phenotype switching in part by enhancing p21 levels (Carreira et al., 2005), we used immunofluorescence to investigate the correlations between BRD inhibition and p21 levels in A375 cells (**Figure 3D**). Consistent with their effects on A375 cell morphology, JQ1 and ABBV-075 treatment led to increased p21 levels (**Figure 3E**). These results were reproducible in melanoma cell lines derived from primary (IGR37) and metastatic (IGR39) sites in a single patient (**Figure S4C-D**). Interestingly, ABBV-744, and to a lower degree the BRD7/9 inhibitor TP-472, induced p21 expression without morphological changes despite limited changes to the BRD4 interaction network compared to those induced by the pan-BET inhibitor ABBV-075 (**Figure S4E**). Our results reveal that BET BRD inhibitors have distinct and much more pronounced effects on melanoma than the other BRD-containing proteins investigated, underscoring their promise as targets for its clinical intervention.

### The MK2 kinase contributes to BET inhibitor sensitivity in melanoma cells

To further elucidate the mechanisms by which BET inhibition impacts cells, we performed a CRISPR/Cas9-based resistance screen for JQ1 in A375 melanoma cells (**Figure S5**). Loss of either p53 or p21 promoted JQ1 resistance (**Figure 4A**), consistent with previous reports showing significant accumulation of cyclin-dependent kinase inhibitors (p21 and CDKN1B (p27)) and cell cycle arrest following BET inhibition (Segura et al., 2013). Our screen also revealed a novel role in JQ1 resistance for MAPKAPK2 (MK2; **Figure 4A**), a kinase downstream of the p38 stress signaling pathway that regulates cell cycle control, chromatin remodeling, UV light-induced DNA damage, and RNA binding (Borisova et al., 2018). To investigate this novel role, we first created a clonal *MK2* knockout (KO) line in A375 cells (*MK2*Δ), which confirmed that loss of MK2 enhanced resistance to BET inhibition (**Figure 4B**). Western blots further revealed that BET inhibition in these lines led to decreased MYC levels with a concomitant increase in p21 (**Figure 4C**). Moreover, the levels of known MK2 substrates were altered with the levels of HSPB1 (HSP27) and its phosphorylated proteoform being reduced while CDC25B was increased over a 48 h JQ1 time course (**Figure 4C**). While analyzing previously published chromatin immunoprecipitation sequencing (ChIP-Seq) datasets in SK-MEL-147 melanoma cells (Vardabasso et al., 2015), we observed that BRD2 localized to the *HSPB1* promoter in a BET inhibition-sensitive manner. We thus investigated how *MK2* KO impacted the transcriptional regulation of *HSPB1* by BRD2 and BRD4 in A375 cells. Using ChIP coupled to quantitative PCR (ChIP-qPCR), we found that JQ1 reduced BRD2 binding to the *HSPB1* promoter and that *MK2* KO significantly reduced this effect (**Figure 4D**); however, it did not rescue a similar decrease in BRD4 binding (**Figure 4E**). To determine if these JQ1-induced binding changes altered HSPB1 and CDC25B expression in MK2 -dependent manner, we measured their nascent RNA levels (**Figure 4F-4G**). After a 24 h JQ1 treatment, the decrease in HSPB1 mRNA was not as pronounced upon *MK2* KO (**Figure 4F**), in line with the protein levels we quantified. This effect was even more pronounced for nascent CDC25B mRNA, which was significantly increased in our *MK2*Δ line (**Figure 4G**), consistent with altered cell cycle regulation (**Figure 4H**). The increased CDC25B levels observed by western blotting are also in agreement with reduced proximity interactions (mapped by TurboID of CDC25B) with ubiquitin-related proteins, including E3 ligases in the APC/C and the substrate recognition protein FBXW11 upon prolonged JQ1 treatment (**Table S2I**). Based on these findings, we hypothesized that BET proteins are also MK2 phosphorylation substrates and that the MK2-dependent phospho-BET proteoforms contribute to HSPB1 and CDC25B transcriptional regulation. To test this possibility, we employed three orthogonal approaches namely: MK2 radioactive kinase assays employing full-length BRD2 or BRD4 purified from K562 cells (*data not shown*); MK2 kinase assays on peptide SPOT arrays containing all putative MK2 phosphorylation motifs (R-X-X-S) and previously reported phosphorylation sites in BRD2, BRD3, and BRD4 (**Figure S6A**; **Table S2J**); and, phosphoproteomics analysis of parental A375 or *MK2*Δ cells treated with and without BET inhibition (**Table S2K**). All three assays failed to detect a direct modulation of BET phosphorylation by MK2 suggesting that MK2 impact on BET BRD inhibitor resistance is through indirect mechanisms.

**Figure 4.**
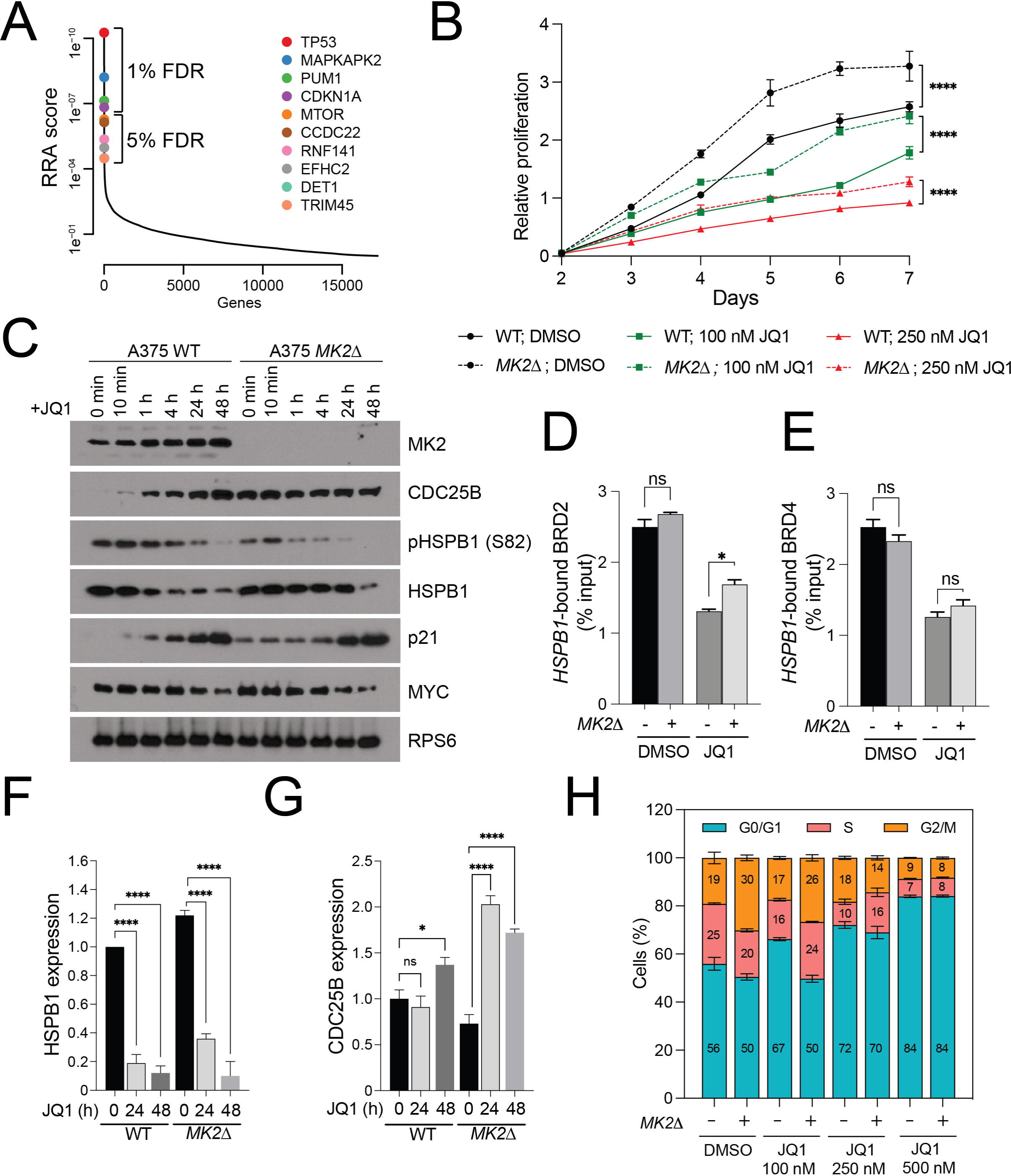
– MK2 is a novel BET inhibitor resistance gene. (**A**) The top 10 hits from the CRISPR/Cas9 JQ1 resistance screen with their associated false discovery rates (% FDR) and robust rank aggregation (RRA) scores. (**B**) Proliferation of parental and *MK2*Δ A375 cells treated with DMSO or 100 nM or 250 nM JQ1 for 7 d. (**C**) Western blot analyses of the indicated proteins following a 48-h time course with 500 nM JQ1 in parental and *MK2*Δ A375 cells. RPS6 was used as a loading control. (**D**-**E**) ChIP-qPCR analyses of BRD2 (**D**) and BRD4 (**E**) on the *HSPB1* promoter in WT or *MK2*Δ A375 cells treated with and without 500 nM JQ1 for 48 h. (**F**-**G**) Nascent-qPCR analyses of HSPB1 (**F**), and CDC25B (**G**) in WT or *MK2*Δ A375 cells treated with and without 500 nM JQ1 for 48 h. Expression levels were normalized to GAPDH. (**H**) Cell cycle analysis of WT or *MK2*Δ A375 cells treated with DMSO or 100, 250, or 500 nM JQ1 for 48 h.

### Inhibiting p38 partially antagonizes BET inhibitors

As loss of *MK2* enhanced the resistance of melanoma cells to BET inhibitors, we next examined the signaling pathways activating MK2 in these cells. To do so, we purified endogenous MK2 from A375 cells treated with (+)-JQ1 or its inactive isomer (-)-JQ1 and analyzed the purified material by mass spectrometry. MK2 interacted with two isoforms of p38, MAPK11 and MAPK14, and many other kinases, suggesting a role as a signaling hub downstream of p38 in melanoma cells (**Figure 5A; Table S2L**). As previously reported (Landry et al., 1992), a potent inhibitor of the p38 MAPK11 and MAPK14 isoforms (SB203580) reduced HSPB1 phosphorylation at S82 (**Figure 5B**), a proxy for MK2 activation. Co-treating A375 cells with SB203580 and JQ1 drastically reduced p21 and CDC25B induction and prevented the loss of HSPB1 expression (**Figure 5B**). Intriguingly, MK2 accumulation following JQ1 treatment was not abolished by SB203580 (**Figure 5B**). Like *MK2* KO (**Figure 4B**), SB203580 treatment enhanced the proliferation of parental A375 cells (**Figure 5C**). Furthermore, A375 cells treated with low concentrations of JQ1 and SB203580 allowed for greater proliferation than in cells treated with JQ1 alone (**Figure 5C**). Lastly, co-treatment with JQ1 and SB203580 reduced the extended cell phenotype (**Figure 5D**) and the proportion of p21-positive cells (**Figure 5E**). Next, we investigated the link between JQ1 and SB203580 across a panel of melanoma cell lines. As in A375 cells, co-treatment minimized the impact of BET inhibition on p38 pathway activation, as assessed using the phospho-shift in MK2 and pHSPB1 (S82) levels, which prevented the loss of HSPB1 (**Figure S6B-D**). To assess whether high HSPB1 expression impacts melanoma *in vivo*, we ranked the TCGA melanoma cohort by HSPB1 level and calculated scores for each tumor representing the expression levels of proliferative (**Figure S6E**) and invasive gene expression signatures (**Figure S6F**) defined by Verfaillie *et al*. (Verfaillie et al., 2015). HSBP1 levels were significantly correlated with the proliferative gene signature (**Figure S6E**), consistent with our *in vitro* observations. Therefore, we propose that inhibiting p38 signaling partially antagonizes BET inhibitors to sustain melanoma proliferation.

**Figure 5.**
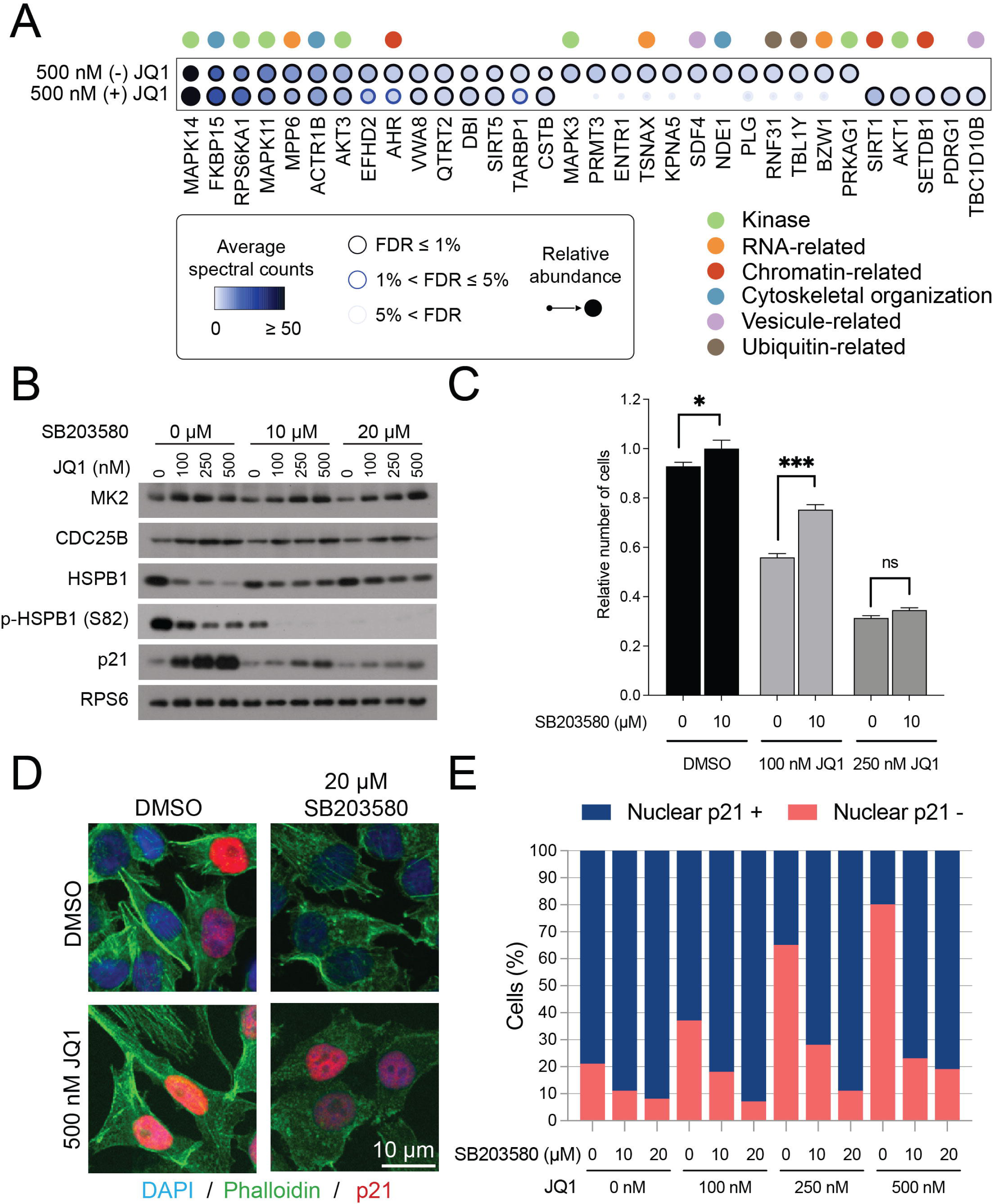
– The p38 signaling pathway partially antagonizes BET BRD inhibitors. (**A**) Dot plot of selected interaction partners of endogenous MK2 from A375 cells treated with 500 nM (+) JQ1 or (-) JQ1 for 48 h. (**B**) Western blot analyses of the indicated proteins in A375 cells following 48-h treatments with the indicated compounds. (**C**) Proliferation of A375 cells treated for 48 h with the indicated compounds. Cell numbers are shown relative to the 10 µM SB203580/DMSO-treated samples. (**D**) A375 cells were treated with the indicated compounds for 48 h and stained with anti-p21 antibodies, phalloidin, and DAPI. (**E**) Quantification of cells treated as in **D** displaying nuclear p21 immunofluorescence.

### BET inhibition locally and globally reorganizes the transcriptional machinery to impact cellular metabolism

To define the molecular mechanisms by which p38 inhibition antagonizes BET inhibitors and sustains higher transcription levels for key transcripts (*e.g*., HSPB1, MYC), we next investigated the transcriptional machinery functionally and physically linked with BET proteins on chromatin (Kim et al., 2019; Lambert et al., 2019; Shu et al., 2020). Specifically, we focused on the DRB sensitivity-inducing factor (DSIF) and negative elongation factor (NELF) complexes and RNA polymerase II (RNAPII) itself, which all actively help transition RNAPII from a promoter-proximal paused state to a productive elongation state. Using ChIP-qPCR, we observed that BRD2 and BRD4 localization to the *MYC* promoter were increased in the *MK2*Δ line, while those of SPT4 (a DSIF subunit), NELFB (a NELF subunit), and POLR2A (RNAPII subunit) were reduced (**Figure 6A**). After a 48-h JQ1 treatment, their levels on the *MYC* promoter were drastically reduced in parental cells compared to DMSO treatment; this effect was significantly minimized by *MK2* KO (**Figure 6A**). This suggest that in the absence of *MK2*, the impact of JQ1 on MYC transcription is not as pronounced as in the parental cell line allowing for more higher MYC levels (**Figure 4C**). To obtain a global view of the transcriptional machinery, we quantified NELF complex foci, which mark sites of intense transcription (Narita et al., 2007). In parental cells, we observed a significant decreased in NELF foci after JQ1 treatment (**Figure 6B-C**). *MK2* KO increased NELF foci numbers in DMSO-treated cells and minimized their loss due to JQ1, further supporting the retention of the transcriptional machinery at sites of active transcription in the *MK2*Δ line upon BET inhibition (**Figure 6B-C**). We confirmed that this was a global phenomenon for BRD2 and BRD4 using CUT&RUN (**Figure 6D-E; Figure S7A-C**). This model is also consistent with the top 100 cancer cell co-dependencies (queried through the DepMap portal (McFarland et al., 2018)) for *BRD2*, and to a lower degree for *BRD4*, which were highly correlated with those of TFIID and NELF complex subunits (**Figure 6F**).

**Figure 6.**
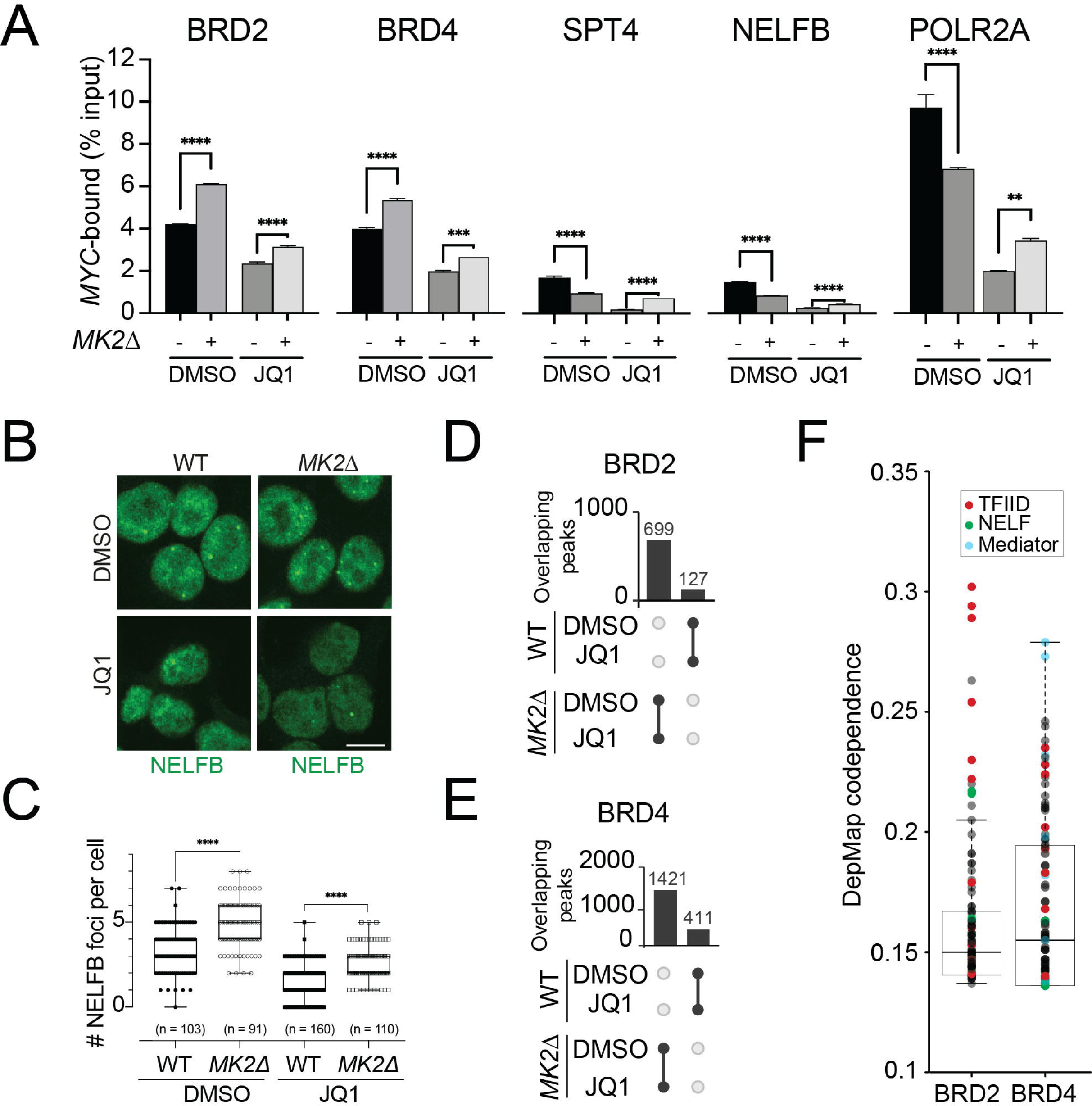
– Loss of MK2 promotes retention of the transcriptional machinery on the chromatin. (**A**) ChIP-qPCR analysis of BRD2, BRD4, SPT4 (DISF complex), NELFB (NELF complex), and POLR2A (RNAPII complex) binding to the *MYC* promoter in WT or *MK2*Δ A375 cells treated with or without 500 nM JQ1 for 48 h. (**B**) A375 cells were treated with the indicated compounds for 48 h and immunostained for NELFB. (**C**) Quantification of NELFB foci in cells treated as in **B**. (**D–E**) UpSet plots of BRD2 (**D**) and BRD4 (**E**) CUT&RUN data from WT or *MK2*Δ A375 cells treated with or without 500 nM JQ1 for 48 h. Only peaks identified in a given cell line +/− JQ1 are shown. (**F**) Whisker plot of positive co-dependencies (Pearson correlations) for BRD2 and BRD4 identified by the DepMap consortium.

Having established that *MK2* KO reorganizes the transcriptional machinery upon BET inhibition, we performed RNA-Seq to identify the most affected transcripts (**Figure 7A-C; S7E**). Gene set enrichment analysis revealed that metabolism-related processes, notably glycolysis and oxidative phosphorylation (OXPHOS), were among those most significantly upregulated by JQ1 in A375 cells (**Figure 7D**). As *MK2* KO reduced the enrichment of glycolysis- and OXPHOS-related transcripts upon BET inhibition (**Figure 7D-F**), we measured the rates of oxygen consumption, mitochondrial respiration, and extracellular acidification (a proxy for aerobic glycolysis) to elucidate MK2’s role in regulating energy metabolism. We observed a significant increase in basal respiration upon *MK2* loss (**Figure 7G-I**). While basal respiration was not significantly altered by 48 h of BET inhibition in WT cells, JQ1 significantly decreased it in *MK2*Δ cells (**Figure 7G-H)**. Furthermore, *MK2*Δ cells had significantly lower maximum respiration and spare respiratory capacities compared to WT cells (**Figure 7J-K**). These results suggest that *MK2* loss induces mitochondrial stress, and related gene signatures are enriched as a stress response mechanism. However, despite enrichment in the OXPHOS gene signature, *MK2*Δ cells exhibited lower mitochondrial capacities. Conversely, basal glycolysis was significantly decreased by 48 h of JQ1 treatment in parental A375 cells but not in *MK2*Δ cells (**Figure 7L**). These alterations suggest a metabolic reprogramming and increased formation of reactive oxygen species upon JQ1 treatment which are alleviated, in part, through the inhibition of the p38-MK2 signaling pathway to sustain proliferation (**Figure 7M**).

**Figure 7.**
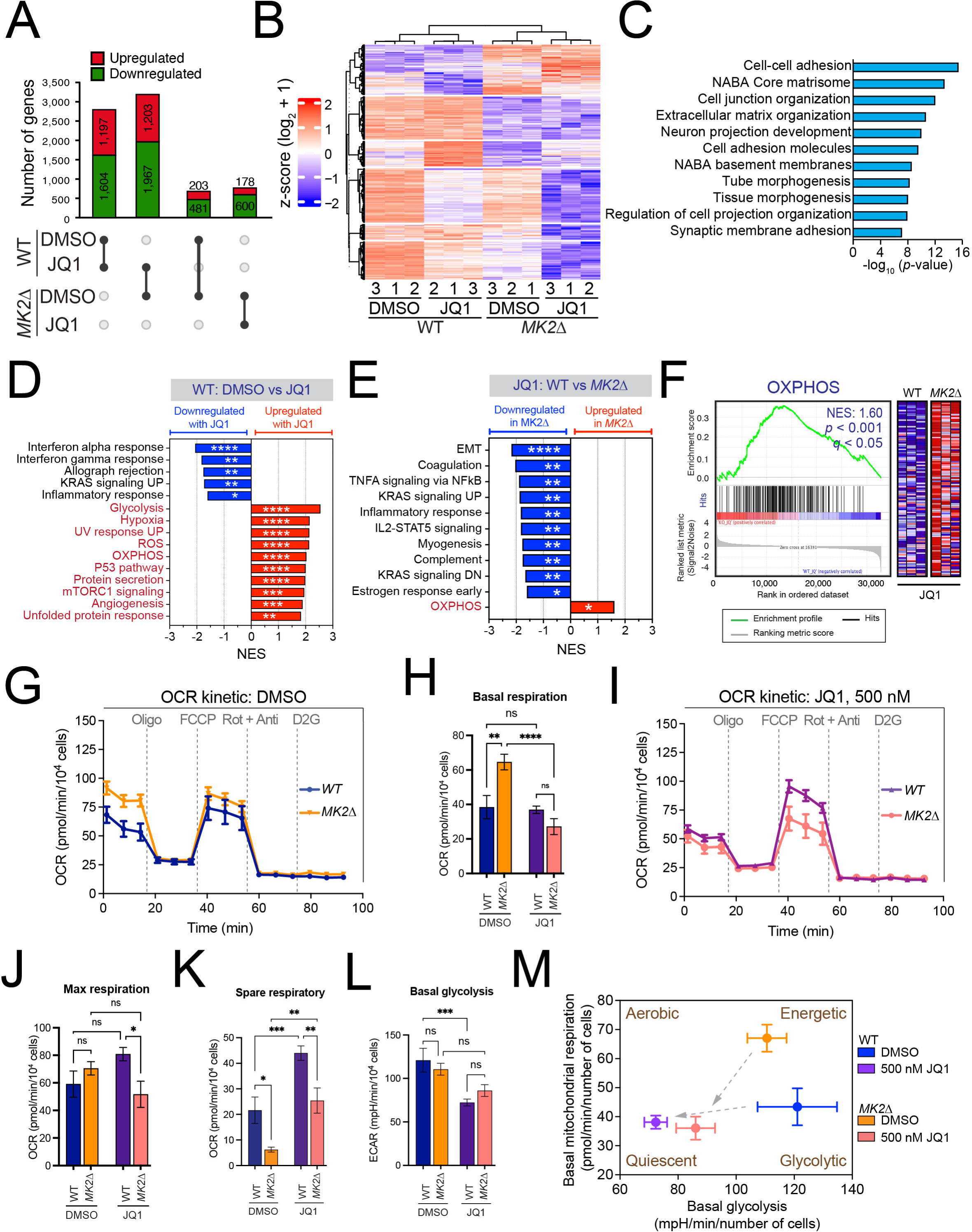
– Loss of MK2 alters the metabolic response of melanoma cells to BET BRD inhibitors. (**A**) Bar graph showing the numbers and proportions of differentially expressed genes between the indicated sample pairs. (**B**) Heat map of the differentially expressed transcripts. (**C**) Enriched GO terms among JQ1-dependent differentially expressed genes. (**D**–**E**) Gene set enrichment analyses of RNA-Seq data from WT A375 cells treated with or without 500 nM JQ1 for 48 h (**D**) and WT and *MK2*ΔA375 cells treated with 500 nM JQ1 for 48 h (**E**). *, ** and **** indicate *p* < 0.05, 0.01, and 0.0001, respectively, by Student’s *t*-test. (**F**) Gene set enrichment analysis plots of changes in the OXPHOS gene signatures in WT or *MK2*Δ A375 cells treated with 500 nM JQ1 for 48 h. The normalized enrichment score (NES), *p*-value, and *q*-value are indicated and only core genes are shown. (**G**) Oxygen consumption rates in DMSO-treated WT or *MK2*Δ A375 cells, normalized to the number of cells. (**H**) Bar graph of the basal respiration of DMSO-treated WT or *MK2*Δ A375 cells (derived from **G)**. ** and **** indicate *p* < 0.01 and 0.0001, respectively, by one-way analysis of variance (ANOVA). (**I**) Oxygen consumption rates in WT or *MK2*Δ A375 cells treated with 500 nM JQ1 for 48 h, normalized to the number of cells. (**J**–**K**) Bar graphs of the maximal respiration rate (**J**) and spare respiratory capacity (**K**) of WT or *MK2*Δ A375 cells treated with 500 nM JQ1 for 48 h (both derived from **I**). ** and **** indicate *p* < 0.01 and 0.0001, respectively, by one-way ANOVA. (**L**) Bar graph of the basal glycolysis rates of WT and *MK2*Δ A375 cells treated with 500 nM JQ1 or DMSO for 48 h. ***, *p* < 0.001 by one-way ANOVA. **(M)** Overview of a Seahorse mitochondrial stress test measuring the oxygen consumption and extracellular acidification rates of WT or *MK2*Δ A375 cells treated with 500 nM JQ1 or DMSO for 48 h.

## Discussion

In this study, we defined the interactomes of BRD-containing proteins and selected functional and physical interactors to shed light on the Kac recognition process. Besides providing a wealth of data for the biological investigation of specific BRD-containing proteins in future studies, our results confirm the predicted concentration of acetylation readers in the nucleus, segregated from the distinct mitochondrial and cytoplasmic metabolic environments (Trefely et al., 2022). The interactome was enriched for known Kac-modified proteins (Hornbeck et al., 2019), and, as first noted for kinases (Breitkreutz et al., 2010; Varjosalo et al., 2013), revealed extensive interconnections between Kac machinery components that could impart specificity in this signaling network.

Our data allowed us to explore the functions of BRD-containing proteins as modular scaffolds (Pawson and Scott, 1997) by mapping most BRD-containing proteins *via* orthogonal proteomic approaches. We generated rich and partially overlapping static interactomes by AP-MS and BioID, further reinforcing their complimentary nature (Lambert et al., 2015). We also investigated how BRD inhibitors remodeled these interactions. In contrast to our previous findings with BET proteins (Lambert et al., 2019), BRD inhibitors targeting CREBBP/EP300 and mSWI/SNF subunits had limited impact on their interactomes. While these results were consistent with the displacement of BRD-containing proteins upon chromatin hypoacetylation (Loehr et al., 2022), this suggests that the ability of BRD inhibitors to outcompete with BRD-containing proteins for chromatin-associated proteins may be limited to a subset of targets in which BRDs are the main reader modules. This is consistent with the overall weak impacts of BRD inhibitors as monotherapies in clinical trials (*e.g*., (Cescon et al., 2024; Hamilton et al., 2023)).

We identified the MK2 kinase as a new BET inhibition resistance gene in melanoma that significantly regulates glycolysis and OXPHOS. This is reminiscent of a report by Santamans *et al*., which showed that p38 kinases, especially MAPK12, were required for metabolic adaptation during physiological exercise (Santamans et al., 2021). Additionally, MAPK12 overexpression leads to glucose intolerance and insulin resistance, whereas early postnatal cardiac-specific *MAPK12*/*MAPK13* deletion increases cardiac glycogen storage and affects whole-body metabolism (Santamans et al., 2021). Mechanistically, loss of MK2 activity through genomic ablation or p38 inhibition retained the transcriptional machinery and BET proteins at pro-proliferative loci, despite concomitant treatment with BET inhibitors. These results are consistent with our observations that *MK2* loss enables the establishment of a transcriptional program favoring OSPHOS, resulting in enhanced basal and maximal respiration. However, loss of MK2 activity in the context of prolonged JQ1 treatment reduces maximum respiration and spare respiratory capacity. This difference may be exploitable, as acute myelogenous leukemia cells with low spare capacity are particularly susceptible to oxidative stress (*e.g*., (Sriskanthadevan et al., 2015)).

While most BET inhibitors developed to date have effectively engaged both BRDs (*i.e*., BD1 and BD2), the recent disclosure of BD1-specific (*e.g*., GSK778 (Gilan et al., 2020)) and BD2-specific (*e.g*., RVX-208 (Picaud et al., 2013), ABV-744 (Faivre et al., 2020), GSK046 (Gilan et al., 2020)) inhibitors has allowed interrogation of their individual functions. BD1-specific inhibitors generally mimic pan-BET inhibitors, while BD2-specific compounds have more subdued impacts (Filippakopoulos and Knapp, 2020). Our quantitative mapping of the BRD4 interaction network with both pan-BET inhibitors and BD2-specific inhibitors further supported these observations. Specifically, changes in the BRD4 interaction network induced by the BD2-specific ABBV-744 had lower amplitudes than those of the pan-BET inhibitors tested (JQ1 and ABBV-075; **Figure 3B**). ABBV-744 treatment in A375 cells resulted in clear p21 induction but did not display reduced MYC levels or the extended cellular phenotype (**Figure 3C-D**). This is consistent with a recent publication showing that the BD1-specific GSK778, but not the BD2-specific GSK046, displaced BETs from the *MYC* enhancer and significantly reduced its transcript level in acute monocytic leukemia cells (Gilan et al., 2020). In contrast to JQ1, which caused BRD4 to localize to the centromere *via* normally weak or neomorphic associations with CENPB and associated factors (**Figure 3B**), ABBV-075 and ABBV-744 did not induce any abundant novel interaction partners. Moreover, both had similar impacts in terms of overall modulation (**Figure S4E**), suggesting that they function through mechanisms that phenocopy each other. As such, our findings support the general notion that the use of BD-specific BET inhibitors may allow for a “separation of functions” that may lead to improved outcomes in clinical trials.

Our results show that p38/MK2 signaling contributes to BET inhibition resistance in the context of cutaneous melanoma exhibiting hyperactive RAS/MAPK signaling. As RAS/MAPK and p38 signaling are highly interconnected (*e.g*., (Ludwig et al., 1998; Zaru et al., 2007)), it is unknown whether our findings will be transposable to other cancer contexts with more controlled RAS/MAPK signaling. As we were unable to identify the MK2 phosphorylation target(s) whose lack of phosphorylation contributes to BET inhibition resistance despite the use of three orthogonal assays, we favor a model in which MK2 regulates BET indirectly. More specifically, we postulate that MK2-dependent phosphorylation of unidentified substrate(s) contributes to BET inhibition resistance by favoring the retention of BET-associated transcriptional machinery at key regulatory regions. As such, the identification of the MK2 substrates contributing to BET inhibitor resistance may represent attractive targets for combination therapy with BET inhibitors.

This study demonstrates how proteomics approaches, coupled with selective inhibition of Kac-dependent readouts, can inform on the intricate connectivity of the acetylation circuitry, while offering insights into novel biological functions for its components. The continued development of selective BRD inhibitors provides opportunities to expand our understanding of Kac signaling while establishing a rationale for translational research that potentiates current treatments targeting it in disease.

## Author contributions

Conceptualization, J.P.L Methodology, J.P.L and A.C.G.; Software, J.D.R.K. Investigation, P.E.K.T., J.L., Z.S., M.T., L.G., A.L. and J.P.L.; Formal Analysis, P.E.K.T., L.G., J.P.L. Resources, S.P., B.G.B., C. G.; Data Curation, J.P.L.; Writing – Original draft, E.A.W., P.F., A.C.G. and J.P.L.; Writing – Review and Editing, E.A.W., P.F., A.C.G. and J.P.L.; Supervision, S.A., E.A.W., P.F., A.C.G and J.P.L.; Project Administration, A.C.G and J.P.L.; and Funding Acquisition, P.F., A.C.G. and J.P.L.

## Acknowledgments

We thank Pavel Savitsky for his help with plasmids generation and the following investigators for the kind gift of reagents: Kyle M. Miller (ATAD2, PBRM1, SP100, SP110, and SP140 cDNAs), Patrick D. Varga-Weisz (BAZ1A cDNA), Joseph W. Landry (BPTF cDNA), Misao Ohki (KMT2A cDNA), David C. Page (TAF1L cDNA), Konstantin Khetchoumian (TRIM66 cDNA), and Jacques Côté (KAT5 cDNA).

---

**Supplemental Figure S1.** – *Expression of the Kac machinery in HEK293 cells and reproducibility metrics* (**A**) Transcript and protein expression levels of all human BRD-containing proteins (grouped by BRD family) (Filippakopoulos et al., 2012) in HEK293 cells. RNA-Seq levels were obtained from the Human Protein Atlas (Uhlen et al., 2015) and Ensembl version 78.38 and are expressed as fragments per kilobase of transcript per million mapped reads (FPKM). Protein levels were obtained from (Schaab et al., 2012) and are expressed as intensity-based absolute quantification (iBAQ) values. (**B**) Transcript and protein expression levels of all human acetyltransferases and deacetylases in HEK293 cells (see description in **A** for details). GNAT: Gcn5-related N-acetyltransferases; CBP: CREB-binding proteins; MYST: MOZ, YBF2, SAS2 and Tip60; NRC: nuclear receptor coactivator. (**C**–**D**) Reproducibility of the spectral counts of high-confidence preys between the two biological replicates. Values are plotted separately for the AP-MS (**C**) and BioID (**D**) datasets. (**E**) Numbers of interactions detected by AP-MS and BioID for the BRD-containing and BRD-associated baits profiled.

**Supplemental Figure S2.** – *Detailed mapping of* cluster 5 *interconnections with the human Kac machinery* (**A**) An interaction network of the high-confidence (FDR ≤ 1%) protein-protein interactions detected for ISWI family members by AP-MS and BioID. (**B**) Violin plots of cancer cell dependencies (Chronos scores) for BRD-containing ISWI family members generated from the 1,100 cell lines screened by the DepMap consortium as of 2023Q4. (**C**) Kaplan-Meier plot of the survival of patients with triple negative breast cancer (TNBC) segregated based on their tumor’s BAZ2B expression level. The plot was generated using the Kaplan-Meier plotter tool (Gyorffy, 2024) and the pan-cancer atlas dataset (Hoadley et al., 2018). (**D**) *Left*, heat map of significant proximity partners of the indicated BAZ2B constructs. *Right*, heat map of the similarities between the BioID profiles of indicated BAZ2B constructs analyzed against the human cell map (humancellmap.org; (Go et al., 2021)).

**Supplemental Figure S3.** - *Mild modulation of CREBBP/EP300 and mSWI/SNF complexes by their BRD inhibitors* (**A**) Dot plots summarizing the unique and shared interactions identified for CREBBP and EP300 in Flp-In T-REx HEK293 cells. (**B**) Overview of the domain architecture of CREBBP and EP300 with the targets of the BRD inhibitors I-CBP112 and SGC-CBP30 and the KAT inhibitor A485 indicated. (**C**) Heat map of significant proximal partners of CREBBP and EP300 in Flp-In T-REx HEK293 cells following 24-h treatments with DMSO, I-CBP112, SGC-CBP30, A485, or A486. (**D**) Heat map of significant proximal partners of PBRM1, SMARCA2, and SMARCA4 in Flp-In T-REx HEK293 cells following 24-h treatments with DMSO, PFI3, or JQ1. Bait proteins are indicated in beige.

**Supplemental Figure S4.** – *Differential impacts of BRD inhibitors on melanoma cell phenotypes and p21 activation* (**A**) A375 cells were treated with the indicated BRD inhibitors for 48 h and stained with anti-p21 antibodies, phalloidin, and DAPI. (**B–C**) IGR37 (**B**) and (**C**) IGR39 cells were treated with (+)-JQ1 or (-)-JQ1 for 48 h and stained with anti-p21 antibodies, phalloidin, and DAPI. (**D**) Bait versus bait scatter plot of BRD4 AP-MS samples from Flp-In T-REx HEK293 cells treated with 500 nM ABBV-075 or 1 µM ABBV-744 for 1 h.

**Supplemental Figure S5.** – *A CRISPR/Cas9-based JQ1 resistance screen in melanoma cells* (**A**) Overview of the CRISPR/Cas9 screen. (**B–E**) Enrichment in individual sgRNAs targeting *TP53* (**B**), *MAPKAPK2* (**C**), *PUM1* (**D**), and *CDKN1A* (**E**). (**F**) Violin plots of cancer cell dependencies (Chronos scores) for JQ1 resistance screen hits based on the 1,100 cell lines screened by the DepMap consortium as of 2023Q4.

**Supplemental Figure S6.** – *Inhibiting p38 signaling antagonizes BET inhibition in melanoma by supporting higher HSPB1 levels* (**A**) MK2 kinase assays on SPOT arrays corresponding to BRD2, BRD3 and BRD4 phosphorylation sites. For each site, the WT peptide sequences and corresponding non-phosphorylatable Ala counterparts (on the central Ser/Thr and on all other phosphorylatable residues, respectively) were synthesized beside it. Peptide sequences are provided in **Table S2J**. (**B–D**) Western blot analyses of the indicated proteins in IGR39 (**B**), SK-MEL-28 (**C**), and SK-MEL-30 (**D**) cells following a 1-h pre-treatment with SB203580 and subsequent JQ1 treatment. (**E–F**) Correlations between HSPB1 levels and gene signatures of (**E**) proliferation and (**F**) invasion from (Verfaillie et al., 2015). The 471 melanomas in the TCGA cohort were ranked by HSPB1 expression (black line). The vertical grey lines indicate HSPB1 expression in each of the tumors, and the red line represents the moving average of the gene signature over each tumor.

**Supplemental Figure S7.** – *Overview of MK2 contribution to BRD2 and BRD4 genomic localization and activity* (**A–C**) Genome browser views of the CUT&RUN signals of BRD2 and BRD4 on *MYC* (**A**), BRD4 on *CUL4B* (**B**), and BRD4 on *ITGA9* (**C**) in parental or *MK2*Δ A375 cells treated with and without 500 nM JQ1 for 48 h. (**D**) Principal component analysis plot of the RNA-Seq results obtained for three biological replicates of each treatment.

**Supplemental Table S1** – *Clone and reagent information*

**Supplemental Table S2** – *AP-MS, BioID, and molecular biology datasets*

