## Supplementaty Methods for "Systematic profiling of the acetyl lysine machinery reveals a role for MAPKAPK2 in bromodomain inhibitor resistance"

**STAR+METHODS:**

Detailed methods are provided in the online version of this paper and include the following:

- **KEY RESOURCES TABLE**
- **CONTACT FOR REAGENT AND RESOURCE SHARING**

- **METHOD DETAILS**
  - Cloning, mutagenesis, and cell line generation
  - FLAG affinity purification
  - Proximity biotinylation
  - Experimental design for mass spectrometry experiments
  - HPLC column preparation for mass spectrometry
  - MS acquisition using an Orbitrap Fusion
  - MS acquisition using an LTQ-Orbitrap
  - Data-dependent acquisition MS analysis
  - MS data visualization and archiving
  - Validation of interactions by immunoblotting
  - Interactome and Kac literature overlap analysis
  - Protein expression and purification
  - Cell cycle analysis
  - ChIP-qPCR and quantitation
  - CUT&RUN
  - RNA-Seq
- **QUANTIFICATION AND STATISTICAL ANALYSIS**
  - CUT&RUN data analysis
  - CUT&RUN peak calling
  - CUT&RUN peak overlap
  - CUT&RUN promoter enrichment
  - CUT&RUN signal density heatmap
  - RNA-Seq differential gene expression (DEG) analysis
  - RNA-Seq gene set enrichment analysis (GSEA)
  - RNA-Seq gene ontology (Metascape) analysis
- **DATA AND SOFTWARE AVAILABILITY**

The MassIVE accession numbers for the MS data reported in this paper are: MSV000080989, MSV000080987, MSV000080996, MSV000080999, MSV000093290, MSV000093292, MSV000093295, MSV000093308, MSV000093309, MSV000093289 and MSV000094608, available at http:// massive.ucsd.edu. Additional files include the complete SAINTexpress outputs for each dataset as well as a ‘‘README’’ file that describes the dataset composition and the experimental procedures associated with each accession number. CUT&RUN and RNA-Seq data reported in this paper were deposited in the Gene Expression Omnibus and assigned the identifiers GSE268319 (reviewer token is “mhwrcguktnwdzgl”) and GSE268320 (reviewer token is “wvkhwwoujvortmj”), respectively.

**METHOD DETAILS**

**Cloning, mutagenesis, and cell line generation**

Flp-In T-REx HEK293, A375 and IGR39 cells were cultured in DMEM high glucose, pyruvate medium supplemented with 10% Fetal Bovine Serum (FBS) and 0.5 μg/mL puromycin. IGR37 cells were cultured in DMEM high glucose, pyruvate medium supplemented with 15% FBS and 0.5 μg/mL puromycin. Constructs for the genes of interest were generated via Gateway cloning into pDEST 5′ 3×FLAG-pcDNA5-FRT-TO or pDEST 5′ BirA*-FLAG-pcDNA5-FRT-TO. Details on all entry clones and destination vectors used in this study can be found in **Tables S1A­–B**. Baits were stably expressed in T-REx Flp-In HEK293 cells as described (Lambert et al., 2014). Parental Flp-In T-REx HEK293 cells and stable cells expressing BirA*-FLAG fused either to green fluorescent protein (GFP) or a nuclear localization sequence (NLS) were processed in parallel to the bait proteins and used as negative controls for the BioID experiments. Flp-In T-REx HEK293 cells expressing NLS-BirA* fused to a FLAG tag were were processed in parallel and used as negative controls for AP-MS experiments. Stable cell lines were grown in the presence of 200 μg/mL hygromycin to 80% confluence before expression was induced *via* 1 μg/mL tetracycline for 24 h (unless otherwise indicated). For BioID experiments, two 150-mm plates were induced with tetracycline and treated with 50 μM biotin for 24 h before harvesting. The cells were harvested and pelleted at low speed, washed with ice-cold phosphate-buffered saline (PBS), and frozen at -80°C until purification. K562 cells stably expressing near-physiological levels of 3×FLAG-Twin-Strep-tagged BRD2, BRD4, or MAPKAPK2 from the *AAVS1* safe harbour locus were established as described previously (Dalvai et al., 2015). These cell lines were cultured in RPMI medium supplemented with 10% newborn calf serum (NBCS) and 0.5 μg/mL puromycin.

**FLAG affinity purification using a chromatin-optimized protocol**

To identify interactions for BET proteins that are either occurring on chromatin, the nucleoplasm, or elsewhere, we used the chromatin-optimized FLAG AP-MS protocol from (Lambert et al., 2014) with slight modifications. Essentially, this protocol incorporates DNA shearing by sonication and nucleases to solubilize DNA-associated protein complexes while largely maintaining protein-protein interactions. Stable cells from two 150-mm plates were pelleted, frozen, and lysed in 1.5 mL ice-cold low salt lysis buffer (50 mM HEPES-NaOH pH 8.0, 100 mM KCl, 2 mM ethylenediaminetetraacetic acid (EDTA), 0.1% NP-40, and 10% glycerol, with 1 mM phenylmethylsulfonyl fluoride (PMSF), 1 mM dithiothreitol (DTT), and Protease Inhibitor Cocktail (Sigma-Aldrich, P8340, 1:500) added immediately prior to processing). To aid with lysis, the cells were frozen on dry ice, thawed in a 37°C water bath, and returned to ice. The samples were sonicated with a QSONICA 125W sonicator equipped with a 1/8” probe at 4°C using three 10 s bursts with 2 s pauses at 35% amplitude. Turbonuclease (100 U) was added and the lysates were rotated at 4°C for 1 h. The lysates were centrifuged at 20,817 × *g* for 20 min at 4°C and each supernatant was added to a tube containing 25 μL of a 50% slurry of magnetic anti-FLAG M2 beads (Sigma-Aldrich, M8823) prewashed with low salt lysis buffer. FLAG immunoprecipitation was allowed to proceed at 4°C for 2 h with rotation. The beads were pelleted by centrifugation (1,000 rpm for 1 min) and magnetized, and the unbound lysate was aspirated and kept for analysis. The beads were demagnetized, washed with 1 mL lysis buffer, and remagnetized to aspirate the wash buffer. The beads were then washed with 1 mL of 20 mM Tris-HCl (pH 8.0) containing 2 mM CaCl_2_ and the excess wash buffer was removed by centrifuging the beads, magnetizing them, and pipetting off the remaining liquid. The now-dry magnetic beads were removed from the magnet and resuspended in 7.5 μL of 20 mM Tris-HCl (pH 8.0) containing 750 ng of trypsin (Sigma-Aldrich, T7575) and the mixture was incubated overnight at 37°C with agitation. After the initial incubation, the beads were magnetized and the supernatant was transferred to a fresh tube. Another 250 ng of trypsin was added and the sample was further digested without agitation for 3–4 h. The tryptic digest was acidified with formic acid to a final concentration of 2% and stored at −40°C until MS analysis.

**Proximity-dependent biotinylation (BioID)**

BioID was performed as in (Lambert et al., 2015) with slight modifications. Cells from two 150-mm plates were pelleted, frozen, and thawed in 1.5 mL ice-cold radioimmunoprecipitation (RIPA) buffer (50 mM Tris-HCl (pH 7.5), 150 mM NaCl, 1% NP-40, 1 mM EDTA, 1 mM ethylenebis(oxyethylenenitrilo)tetraacetic acid (EGTA), 0.1% sodium dodecyl sulfate (SDS), and 0.5% sodium deoxycholate) with PMSF (1 mM), DTT (1 mM), and Protease Inhibitor Cocktail (Sigma-Aldrich) added immediately before use). The lysates were sonicated using a QSONICA 125W sonicator equipped with a 1/8” probe, treated with Benzonase, and centrifuged as described in the FLAG AP-MS section. For each sample, 60 μL of streptavidin-sepharose bead slurry (Cytiva, 17511301) was washed three times with 1 mL of RIPA buffer by pelleting the beads *via* gentle centrifugation at 500 × *g* for 1 min and aspirating the supernatant, then added to the lysate to capture biotinylated proteins for 3 h at 4°C with rotation. The beads were gently pelleted and then washed twice with 1 mL RIPA buffer and three times with 1 mL 50 mM ammonium bicarbonate (pH 8.0). After the final wash, the beads were pelleted and any excess liquid was aspirated off. The beads were re-suspended in 100 μL of 50 mM ammonium bicarbonate ~ pH 8.0, and 1 μg of trypsin solution was added. The samples were rotated overnight at 37°C and then an additional 1 μg of trypsin was added, followed by further incubation for 2–4 h. The beads were pelleted and the supernatant was transferred to a fresh tube. The beads were rinsed twice with 100 μL high-performance liquid chromatography (HPLC)-grade water, and the washes were combined with the supernatant. The peptide solution was acidified with 50% formic acid to a final concentration of 2% and the sample was dried in a SpeedVac. Tryptic peptides were resuspended in 25 μL 5% formic acid and stored at -80°C until MS analysis.

**Phosphopeptide analysis**

A375 WT and MK2 KO cells were cultured in complete DMEM medium and treated with 0.5 µM JQ1 or DMSO for 48 hours. Two 15-cm plates were set for each condition. After treatment, the cells were washed with 1xPBS, harvested using a spatula. Cells were pelleted at 4°C, snap frozen on dry ice and stored at -80°C before proteins purification. Cells pellets were thawed on ice and resuspended in 1.5 mL RIPA buffer freshly supplemented with 1 mM PMSF, 1 mM DTT, 1× Protease Inhibitor Cocktail (Sigma-Aldrich, Cat#P8340), Phosphatase Inhibitor Cocktail 2 (Sigma-Aldrich, #P5726), and Phosphatase Inhibitor Cocktail 3 (Sigma-Aldrich, Cat#P0044). Lysates were sonicated using a Sonic Dismembranor Series 60 (four cycles of 10 s on and 20 s off at 35% amplitude), treated with 1 µL Turbonuclease (Sigma; #T4330), and rotated for 1 h at 4°C. Samples were centrifuged at 12,000 × *g* for 20 min at 4°C and the supernatants were recovered into new tubes. Proteins were non-selectively extracted and digested using the SP3 protocol with a few modifications (Hughes et al., 2019; Moggridge et al., 2018). Briefly, 20 mg SeraMag beads (GE Healthcare, 65152105050250 and 45152105050250, mixed at a 1:1 ratio) were washed twice with 1 mL H_2_O and added to 2 mg of protein. The final volume was adjusted to 1 mL with 50% ethanol. Samples were incubated for 20 min at room temperature with horizontal rotation to allow the beads to bind the proteins. After magnetizing the beads and washing them three times with 500 µL of 80% ethanol, the beads were transferred to new tubes, resuspended in 400 µL of 50 mM HEPES pH 8.0 containing 40 µg trypsin (Sigma-Aldrich, Cat#T6567), bath-sonicated for 15 s, and incubated overnight at 37°C with lateral rotation. The resulting peptides were recovered in new tubes, the beads were rinsed twice with 300 µL acetonitrile, which was pooled with the peptides to maximize recovery, and formic acid was added to a final concentration of 2%. An aliquot containing 10% of the peptides was retained as an input sample and 90% were used for phosphopeptide enrichment. Both fractions were vacuum-dried in a SpeedVac, then desalted using HyperSep C_18_ Cartridges (Thermo, Cat# 60108-301) for subsequent phosphopeptide purification following the manufacturer’s instructions (input samples were desalted on homemade C_18_ StageTips, vacuum-dried, and stored at -80°C until MS analysis). The desalted peptides were subjected to TiO_2_-based phosphopeptide enrichment using a High Select Phosphopeptide Enrichment Kit (Thermo, Cat# A32993) following the manufacturer’s instructions. Briefly, the peptides were resuspended in 150 µL of binding/equilibration buffer, then added to an equilibrated TiO_2_ column. After sequentially washing the column with binding/equilibration buffer, wash buffer, and LC-MS grade water, the phosphopeptides were eluted from the TiO_2_ beads with 50 µL phosphopeptide elution buffer, dried in a SpeedVac, and stored at -80°C until MS analysis.

**Experimental design for MS experiments**

For each bait, two biological replicates were independently processed. Each batch of samples processed included negative controls, which were grown in parallel to the bait samples and treated in the same manner. For AP-MS, we used cells expressing a 3×FLAG-GFP tag construct or no bait (*i.e*. an empty cell line). For BioID, we used cells expressing BirA*-FLAG-GFP, BirA*-NLS-FLAG, or no bait. To minimize sample carry-over issues during liquid chromatography, extensive washes were performed between samples (details for each instrument are provided below), and the order of sample acquisition was reversed between replicate batches.

**MS acquisition using an LTQ mass spectrometer**

The entire resuspended sample was bomb-loaded onto a homemade column equilibrated with buffer A (2% acetonitrile, 0.1% formic acid) as per (Lambert et al., 2014). The column was washed off-line for 10 min in buffer A, then placed in-line with a LTQ mass spectrometer equipped with an Agilent 1100 pump with split flow, and a Thermo or Proxeon source. The HPLC delivered an acetonitrile gradient over 120 min (1–5% buffer B (98% acetonitrile, 0.1% formic acid) over 4 min, 5–40% buffer B over 100 min, 40–60% buffer B over 5 min, 60–100% buffer B over 5 min, hold buffer B at 100% for 3 min, then 100–0% B over 2 min). Data-dependent acquisition (DDA) involved one centroid MS scan (mass range 400–2000) followed by MS/MS scans on the five most abundant ions. General parameters were: activation type = CID, isolation width = 3, normalized collision energy = 32, activation Q = 0.25, activation time = 30 ms, wideband activation. DDA parameters were: minimum threshold = 1000, repeat count = 1, repeat duration = 30 s, exclusion size list = 500, exclusion duration = 30 s, exclusion mass width (by mass) = low 1.2, high 1.5.

**MS acquisition using an Orbitrap Fusion mass spectrometer**

Peptide samples were separated by online reversed-phase nanoscale capillary liquid chromatography and analyzed by electrospray MS/MS. The experiments were performed with a Dionex UltiMate 3000 RSLCnano chromatography system (Thermo Fisher Scientific) connected to an Orbitrap Fusion mass spectrometer (Thermo Fisher Scientific) equipped with a nanoelectrospray ion source. Peptides were trapped at 20 μL/min in loading solvent (2% acetonitrile, 0.05% TFA) on an Acclaim 5μm PepMap 300 μ-Precolumns Cartridge Column (Thermo Fisher Scientific) for 5 min. Then, the precolumn was switched online with a laboratory-made 50 cm × 75 μm internal diameter separation column packed with ReproSil-Pur C18-AQ 3-μm resin (Dr. Maisch HPLC) and the peptides were eluted with a linear gradient of 5–40% solvent B (A: 0,1% formic acid, B: 80% acetonitrile, 0.1% formic acid) over 90 min at 300 nL/min. Mass spectra were acquired in DDA mode using Thermo XCalibur software version 3.0.63. Full scan mass spectra (350–1,800 *m/z*) were acquired in the Orbitrap using an AGC target of 4e5, a maximum injection time of 50 ms, and a resolution of 120,000. Internal calibration using lock mass on the *m/z* 445.12003 siloxane ion was used. Each MS scan was followed by MS/MS scans of the most 10 intense ions for a total cycle time of 3 s (top speed mode). The selected ions were isolated using the quadrupole analyzer in 1.6 *m/z* windows and fragmented by higher energy collision-induced dissociation at 35% collision energy. The resulting fragments were detected by the linear ion trap in rapid scan rate with an AGC target of 1e4 and a maximum injection time of 50 ms. Dynamic exclusion of previously fragmented peptides was set for a period of 20 s and a tolerance of 10 ppm.

**Data-dependent acquisition mass spectrometry analysis**

MS data was stored, searched, and analyzed using the ProHits laboratory information management system (Liu et al., 2016). Thermo Fisher Scientific RAW MS files were converted to mzML and mzXML using ProteoWizard (version 3.0.4468 (Kessner et al., 2008)). The mzML and mzXML files were then searched using Mascot (version 2.3.02) and Comet (version 2012.02 rev.0) against the RefSeq database (version 57, January 30th, 2013) acquired from NCBI, which contains 72,482 human and adenovirus sequences supplemented with common contaminants from the Max Planck Institute (http://141.61.102.106:8080/share.cgi?ssid=0f2gfuB) and the Global Proteome Machine (GPM; http://www.thegpm.org/crap/index.html). For files analyzed on the LTQ, charges of +1, +2, and +3 were considered, at mass tolerances of 3 and 0.6 amu for parent ions and fragments, respectively. For files analyzed on the Orbitrap Fusion, charges of +2, +3, and +4 were allowed, the parent mass tolerance was 12 ppm, and the fragment bin tolerance was 0.6 amu. Deamidated asparagine and glutamine and oxidized methionine were allowed as variable modifications. The results from each search engine were analysed through TPP (the Trans-Proteomic Pipeline (version 4.6 OCCUPY rev 3) (Deutsch et al., 2010) *via* the iProphet pipeline (Shteynberg et al., 2011). Two unique peptide ions and a minimum iProphet probability of 0.95 were required for protein identification. SAINTexpress version 3.3(Teo et al., 2014) was used to calculate the statistical probability of each potential protein-protein interaction compared to background contaminants using default parameters. Unless otherwise specified, controls were compressed by half, to a minimum of eight, using the strategy first introduced in (Mellacheruvu et al., 2013).

**MS data visualization and archiving**

Functional enrichment analyses were performed with g:Profiler (Reimand et al., 2016) using default parameters. Dot plots and heat maps were generated using ProHits-viz (prohits-viz.org (Knight et al., 2017)) and Venn diagrams were generated in R (R Core Team, 2017) using the venneuler package (Wilkinson, 2012). Interaction networks were generated using Cytoscape (V3.5.1; (Shannon et al., 2003)), using the edge thickness to reflect each prey’s average spectral counts. Nodes were manually arranged into physical complexes. All MS files used in this study were deposited in MassIVE (http://massive.ucsd.edu). The username and password to access these files until publication is “BET_inhibition”. Additional details (including MassIVE accession numbers and FTP download links) can be found in **Table S2M**.

**Interactome and Kac literature overlap analysis**

We performed custom downloads of all interactions for each bait protein from BioGRID version 4.4.233, released on May 1^st^, 2024 (Oughtred et al., 2019). Bait-prey and prey-bait relationships were both considered in overlap analyses; for BioGRID, only physical interactions were considered, with no other restrictions regarding experimental evidence. The complete Kac database was obtained from the PTMVar dataset of PhosphoSitePlus (Hornbeck et al., 2019) (www.phosphositeplus.org; June 2016 version).

**Protein complex purification from K562 cells**

BRD2, BRD4 and MK2 were individually purified from 3 L of 3×FLAG-Twin-Strep-tagged K562 cells as described in (Lashgari et al., 2019) with modifications. Briefly, nuclear extracts were prepared following standard procedures and precleared on CL6B Sepharose beads. Tagged proteins were immunoprecipitated with anti-FLAG agarose affinity gel (Millipore Sigma; A2220), then eluted with 200 μg/mL 3×FLAG peptide (Millipore Sigma; SAE0194) in 20 mM HEPES pH 7.5, 150 mM KCl, 0.1 mM EDTA, 10% glycerol, 0.1% Tween-20, and 1 mM DTT supplemented with proteases, deacetylases, and phosphatase inhibitors, followed by affinity purification on Strep-Tactin XT 4Flow agarose beads (IBA; 2-5030-002), and elution with 50 mM D-biotin (prepared fresh by mixing a 1:1 ratio of 100 mM D-biotin stock (in 40 mM HEPES pH 7.9, 150 mM KCl) and 20 mM HEPES pH 7.9, 150 mM KCl, 20% glycerol, 0.2% Tween-20, and 2 mM DTT, supplemented with protease inhibitors). Aliquots of the elutions were resolved on NuPAGE 4–12% Bis-Tris gels (Invitrogen) and visualized by silver staining to ensure their quality.

**Immunofluorescence**

A375, IGR37, and IGR39 cells were seeded onto poly-L-lysine-coated coverslips in 12-well plates in complete medium and grown for 24 h with and without chemical inhibitors as indicated, for 48 h. Cells were fixed with 4% paraformaldehyde in phosphate-buffered saline (PBS) for 15 min at room temperature and then stained using the primary and secondary antibodies listed in the Key Resource Table. Image stacks were acquired on a Leica DMI 6000 B inverted microscope with a Yokogawa CSU10 confocal unit at 63×, deconvolved using Volocity (Quorum Technologies), and shown as intensity projections. Images were cropped using Adobe Photoshop. For all quantitatively compared images, identical imaging conditions (including exposure times) were used.

**Immunoblotting**

For western blot analysis, 10–50 μg of protein was resolved by SDS-polyacrylamide gel electrophoresis, transferred to nitrocellulose membranes, and blocked in Tris-buffered saline containing either 5 mg/mL non-fat milk or bovine serum albumin and 1% Tween-20 for 1 h at room temperature. Antibodies and their conditions can be found in the Key Resource Table. Bands were detected on film using the Clarity Western ECL Substrate (Bio-Rad; #1705061). Films were scanned and figures were assembled using Adobe Photoshop and Adobe Illustrator.

**Cell cycle analysis**

Triplicate A375 WT and MK2 KO cells were plated in 10-cm plates with complete DMEM medium and treated with either DMSO or JQ1 for 48 h. After treatment, cells were trypsinized, and 1 million cells from each sample were transferred into 1.5 mL tubes and washed with 1× PBS. Cells were permeabilized by resuspending them in 1 mL of cold 70% ethanol (diluted with 1× PBS) and incubated overnight at -20°C. After centrifugation, cells were resuspended in 570 µL of 1× PBS supplemented with 0.3 mg/mL RNaseA and incubated for 30 min at 37°C. Subsequently, the samples were transferred into FACS tubes and stained with 30 µL of propidium iodide (1 mg/mL) for 30 min at 4°C in the dark. Flow cytometry data were acquired using a BD FACSCelesta Flow Cytometer with BD FACSDiva software version 8.0.1.1 (Becton, Dickinson and Company). Data analysis was performed using FlowJo version 10.7.1 (Becton, Dickinson and Company), acquiring 10,000 events for each sample.

**Chromatin immunoprecipitation (ChIP)-qPCR and quantitation**

Cells were grown in 150-mm dishes and treated for 48 h with JQ1 or dimethyl sulfoxide (DMSO), washed with 1× PBS and crosslinked with 10 mL of 1% formaldehyde (Sigma, #252549) in 1× PBS at room temperature for 15 min with gentle horizontal rotation. Crosslinking was quenched by incubating the cells with 125 mM glycine for 5 min and washing them three times with 15 mL of ice-cold 1× PBS. Cells were scraped in 1× PBS and transferred to tubes, pelleted at 1,000 × *g* for 1 min, resuspended in ice-cold lysis buffer (50 mM Tris-HCl pH 7.5, 1 mM EDTA pH 8.0, 140 mM NaCl, and 1% NP-40, supplemented with Protease Inhibitor Cocktail (Sigma-Aldrich)) at 1 mL/2×10^7^ cells, and incubated on ice for 10 min. The samples were centrifuged at 2,000 × *g* for 10 min at 4°C and each nuclei-containing pellet was resuspended in ice-cold nuclear lysis buffer (50 mM Tris-HCl pH 7.5, 2 mM EDTA pH 8.0, 0.5% sodium deoxycholate, and 1% SDS, supplemented with Protease Inhibitor Cocktail (Sigma-Aldrich)) at 0.5 mL/2×10^7^ cells and sonicated (12 cycles of 30 sec ON/30 sec OFF, high level) at 4°C with a Bioruptor (Diagenode, #B01020001) to generate ~200–500 bp chromatin fragments. The lysate was centrifuged at 20,000 × *g* for 10 min at 4°C, and the cleared soluble chromatin fragments in the supernatant were transferred to a new tube. Most of the sample was flash-frozen and stored at -80°C until ChIP; however, a 20 µL aliquot was retained to determine the DNA concentration and fragment size distribution. To do this, the sheared chromatin aliquot was diluted is diluted with 170 μL dilution buffer and 10 μL of 5 M NaCl and incubate at 65°C overnight. The next day, add 1 μL of 10 mg/mL RNase A (NEB, #T3018L) and incubate for 30 min at 37°C. Then, 5 μL of 20 mg/mL Proteinase K (Invitrogen, #25530049) was added to the reaction mixt and incubated at 55°C for 1 h. The DNA fragments were then purified using a Monarch PCR & DNA Cleanup Kit (New England Biolabs, # T1030S), quantified with NanoDrop (Thermo Scientific), and resolved on agarose gel. ChIP was performed by first diluting the purified DNA fragments in 9 volumes dilution buffer (20 mM Tris-HCl, pH 8.0, 1 mM EDTA pH 8.0, 0.1% sodium deoxycholate, 140 mM NaCl, 0.01% SDS, and 1% NP-40) and incubating them with Dynabeads Protein A beads (Invitrogen, # 10002D) pre-coupled (2.5 µg antibody for 25 µL of beads) to anti-FLAG antibodies (Sigma, #F1804) for 3 h at 4°C with gentle rotation. Rabbit IgG (Sigma, # I8140) was used as negative control. The beads were recovered after immunoprecipitation (IP) and sequentially washed with low-salt buffer (20 mM Tris-HCl pH 8.0, 2 mM EDTA pH 8.0, 0.1% SDS, 1% Triton X-100, 150 mM NaCl), high salt buffer (20 mM Tris-HCl pH 8.0, 2 mM EDTA pH 8.0, 0.1% SDS, 1% Triton X-100, 500 mM NaCl), LiCl buffer (20 mM Tris-HCl pH 8.0, 2 mM EDTA pH 8.0, 0.5% sodium deoxycholate, 0.1% NP-40, 250 mM LiCl), and TE buffer (10 mM Tris-HCl pH 8.0, 1 mM EDTA). IP chromatin fragments were eluted from the beads with the 100 µL elution buffer (1% SDS, 100 mM NaHCO_3_). Samples were crosslink-reversed, treated with RNase A and proteinase K, and purified as described above. Inputs and purified samples were diluted to 50 ng/mL of DNA fragment) and specific DNA loci were quantified by qPCR using Luna Universal qPCR Master Mix (NEB, #M3003) following the manufacturer’s protocol. Reactions were performed in MicroAmp EnduraPlate Optical 96-Well Clear Reaction Plates (Applied Biosystems, Cat# 4483354) on a QuantStudio 3 Real-Time PCR System (Applied Biosystems, Cat#A28567) with SYBR Green chemistry. Three independent experiments per experimental condition were performed and each sample was processed in three technical replicates. The data were analyzed using the fold enrichment method and the results were normalized to glyceraldehyde 3-phosphate dehydrogenase (GAPDH). Primers used for qPCR are listed in **Table S1C**.

**RT-qPCR**

Wild-type (WT) and *MAPKAPK2Δ* A375 cells were treated with and without JQ1, harvested by scrapping in 1× PBS and transferred to tubes, pelleted at 1,000 × *g* for 1 min, washed with 1× PBS, at the indicated timepoints. Their total RNA was immediately extracted using a Monarch Total RNA Miniprep Kit (NEB, Cat#T2010) according to the manufacturer’s protocol. 1 µg of total RNA were immediately reverse-transcribed to cDNA using the SuperScript III First-Strand Synthesis System (Thermo Fisher, Cat#18080051). The cDNA was quantified and diluted, and qPCR was performed with Luna Universal qPCR Master Mix (NEB, Cat# M3003). The samples were run in technical triplicate on a QuantStudio 3 Real-Time PCR System (Applied Biosystems, Cat#A28567) and analyzed using the 2^-ΔΔCt^ method. RT-qPCR was performed on three independent biological replicates, and reactions with abnormal amplification or melting curves were excluded from the analysis. Relative expression was normalized to GAPDH or actin B (ACTB). Primers used for qPCR are listed in **Table S1C**.

**Nascent RNA qPCR**

Newly synthesized RNAs transcripts were purified using the Click-iT Nascent RNA Capture Kit (Invitrogen, Cat# C10365). In this approach, the newly synthesized RNA incorporates 5-ethynyl uridine (EU), an alkyne-modified nucleotide that can be later modified *via* a click reaction with biotin azide. The biotin-labelled nascent transcripts are capturable with streptavidin-functionalized beads, reverse-transcribed to cDNA, and quantified by qPCR. The experiment was performed following the manufacturer’s protocol with slight modifications. Briefly, cells were cultured in 12-well plate with complete DMEM medium for 48 hrs under appropriate treatment. 0.2 mM EU was then added to the medium and the plate was maintained incubated for 4 h under standard culture conditions (37°C, 5% CO_2_, humidified atmosphere) to EU-label the newly synthesized transcripts. The cells were harvested with a scrapper into a 1.5 mL tube and washed with 1× PBS, and the total RNA was purified using a Monarch Total RNA Miniprep Kit. The RNA content was measured with NanoDrop. 5 µg of purified RNA was incubated with 0.5 mM biotin azide in a click reaction cocktail containing CuSO_4_ for 30 min at room temperature with occasional gentle vortexing. To precipitate the RNA, 1 μL of UltraPure Glycogen, 50 μL of 7.5 M ammonium acetate, and 700 μL of chilled 100% ethanol was added to the click reaction mixt and homogenized by gentle vortexing. Samples were then incubated overnight at -80°C, centrifuged at 13,000 g for 20 min at 4°C. The pellet (precipitated RNA) was resuspended in with 700 µL of 75% ethanol, vortexed briefly and centrifuged at 13,000 g for 5 minutes. The dry pellet was then resuspended in 50 µL DNase/RNase-free distilled water, and the RNA content was quantified with NanoDrop. 500 ng of biotinylated total RNA was incubated with 50 µL Dynabeads MyOne Streptavidin T1 magnetic beads for 30 min at room temperature with gentle vortexing. The biotinylated nascent RNA was captured on the beads, washed and immediately reverse transcribed on the beads using a SuperScript VILO cDNA Synthesis Kit (Invitrogen, Cat# 11754050). The synthesized cDNA was subsequently quantified via qPCR using Luna Universal qPCR Master Mix. RT-qPCR was performed on three independent biological replicates and reactions displaying abnormal amplification or melting curves were excluded from the analysis. Relative expression was normalized to GAPDH or ACTB. Primers used for qPCR are listed in **Table S1C.**

**CUT&RUN**

CUT&RUN experiments were performed in triplicate as previously described by (Skene et al., 2018) with modifications. Briefly, WT and *MK2* KO A375 cells were treated with 500 nM JQ1 or DMSO for 48 h, harvested, lysed with StemPro Accutase (Thermo Fisher Scientific catalog no. A1110501) for 5 min, and centrifuged at 300 × *g* for 3 min. Cells (5×10^6^) were cross-linked with 0.1% formaldehyde for 1 min at room temperature, washed twice with ice-cold PBS, and collected by centrifugation at 300 × *g* for 3 min at 4°C.

The nuclei were isolated using nuclear extraction buffer (20 mM HEPES-KOH, pH 7.9, 10 mM KCl, 0.5 mM spermidine (Sigma-Aldrich, catalog no. 05292-1ml-F), 0.1% Triton X-100, 20% glycerol, and 1× Halt Protease and Phosphatase Inhibitor Single-Use Cocktail (Thermo Fisher Scientific, catalog no. 78442)), then bound to 50 μL of activated concanavalin A bead slurry (Bangs Laboratories, catalog no.Bp531) for 15 min at room temperature. The nuclei-bead suspension was then incubated with primary antibodies in antibody buffer (20 mM HEPES-KOH, pH 7.9, 150 mM NaCl, 1% Triton X-100, 0.05% SDS, 0.5 mM spermidine, 1× Halt Protease and Phosphatase Inhibitor Single-Use Cocktail, 0.01% digitonin, and 2 mM EDTA) overnight at 4°C. Unbound antibodies were removed by two washes with wash buffer (20 mM HEPES-KOH, pH 7.9, 150 mM NaCl, 1% Triton X-100, 0.05% SDS, 0.5 mM spermidine, and 1× Halt Protease and Phosphatase Inhibitor Single-Use Cocktail) using a magnetic stand. The supernatants were discarded and CUTANA pAG-MNase for ChIC/CUT&RUN Workflows (EpiCypher catalog no. EP151016) was added at a final dilution of 1:20 and the samples were incubated for 90 min on a nutator at 4°C. The nuclei-bound beads were washed twice with wash buffer and chilled on ice on a metal rack for 5 min before activating the pAG-MNase with 2 mM CaCl_2_ for 30 min still on ice. The digestion reaction was stopped by adding an equal volume of 2× STOP buffer (200 mM NaCl, 20 mM EDTA, 4 mM EGTA, 20 μg/mL RNase A, 50 μg/mL glycogen, and 100 pg/mL of CUTANA *E. coli* Spike-in DNA) followed by incubation for 10 min at 37°C to release the cleaved DNA fragments. The supernatant (containing the targeted protein-DNA complex) was collected *via* centrifugation at 16,000 × *g* for 5 min, and the DNA uncrosslinked overnight by incubating the samples at 55°C. To recover the DNA-protein complexes, the digested DNA was purified using a NEB Monarch PCR and DNA Cleanup Kit (New England Biolabs, catalog no. T1030L) according to the manufacturer’s instructions.

The concentration of the extracted DNA was estimated using Qubit dsDNA Quantification Assay Kits (Thermo Fisher Scientific, Q32851) before library preparation with the NEBNext Ultra II DNA kit for low-input ChIP and 100-bp paired-end sequencing on an Illumina NovaSeq 6000 Sequencing System. Raw reads were trimmed using fastp v0.21.0 (Chen et al., 2018). Trimmed reads were aligned to the human genome (hg38) using bwa mem v0.7.17 (Li, 2013) and SAMtools v1.13 (Li et al., 2009). Raw signal tracks and normalized tracks (in reads per million (RPM)) were produced from mapped reads using deepTools’ v2.17.0 bamCoverage tool (Ramirez et al., 2014) and BEDTools’ genomecov tool (Quinlan and Hall, 2010), respectively. Tracks were converted to the bigwig format using bedGraphToBigWig v2.8 (Kent et al., 2010). MACS2 v2.2.1 software (Feng et al., 2012) was used for peak calling with the following parameters: no λ, fragment size to 14, mfold from 5 to 50, and keeping all duplicated tags. The CUT&RUN files used in the heat map were lifted over to UCSC hg18 using the UCSC lift over tool (Kuhn et al., 2013). Heat maps were generated using the ComplexHeatmap v2.6.2 (Gu et al., 2016) and EnrichedHeatmap v1.20.0 packages (Gu et al., 2018).

The following antibodies were used: rabbit polyclonal anti-BRD4 (Bethyl Laboratories, catalog no. A301-985 A100, lot no. 8), anti-BRD2 (Bethyl Laboratories, catalog no. A302–583, lot no. 6), anti-H3K27ac (Cell Signaling Technology 8173), and anti-Rpb1 CTD(4H8) (Cell Signaling Technology 2629). Rabbit IgG (EpiCypher 13-0042) was used as a negative control antibody.

**RNA-Seq**

RNA from WT and *MK2* KO A375 cells treated with 500 nM JQ1 or DMSO for 48 h was purified using the RNeasy Plus Mini Kit (QIAGEN) according to the manufacturer’s instructions. The integrity of the extracted RNA was confirmed on a TapeStation 2200 (Agilent Technologies, Santa Clara, CA, USA). All the samples had RNA integrity number equivalents ≥ 9.8. The NEBNext Ultra II Directional RNA Library Prep Kit for Illumina (New England Biolabs Inc., Ipswich, MA, USA) was used to prepare RNA-Seq libraries from 1 µg of total RNA according to the manufacturer’s instructions. Cytoplasmic and mitochondrial ribosomal RNA (rRNA) were removed using a NEBNext rRNA depletion kit V2 (New England Biolabs). Following purification with Agencourt RNAClean XP beads (Beckman Coutler, Missisauga, ON, Canada), the RNA was fragmented using divalent cations under elevated temperature, then used as a template for cDNA synthesis by reverse transcriptase with random primers. Strand specificity was obtained by replacing dTTP with dUTP. The cDNA was subsequently converted to double-stranded DNA and end-repaired. Adaptors were ligated to the dsDNA, followed by purification with an AxyPrep Mag PCR Clean-up Kit (Axygen, Big Flats, NY, USA), excision of the dUTP-containing strands, and finally, a nine-cycle PCR enrichment step to incorporate specific indexed adapters for multiplexing. The quality of the final amplified libraries was examined with a DNA ScreenTape D1000 on a TapeStation 2200 (Agilent Technologies) and quantified on a QBit 3.0 fluorometer (Thermo Fisher Scientific). Subsequently, RNA-Seq libraries with unique indices were pooled at equimolar ratios and subjected to 100-bp paired-end sequencing on a NovaSeq 6000 Sequencing System. The average library read size was 300 bp. The mean coverage/sample was 42M paired-end reads.

**CRISPR screen**

Cas9-expressing A375 cells were transduced with the lentiviral TKO library v1 as previously described (Hart et al., 2015) at a multiplicity of infection of ∼0.3, such that every gRNA was represented by > 100 cells. After 24 h, the cells were selected with puromycin for 72 h, then split into four replicates. Two were treated with 500 nM (+)-JQ1; the other two with 500 nM (-)-JQ1 (as an inactive control). The cells were passaged every 3 d and maintained at 100-fold coverage. Cells were collected and pelleted on days 0 and 21 post-selection and the genomic DNA was extracted. The gRNA inserts were amplified by PCR using primers harboring Illumina TruSeq adapters with i5 and i7 barcodes, and the resulting libraries were sequenced on an Illumina HiSeq 2500 Sequencing System as previously described (Hart et al., 2015).

**Seahorse assays**

Mitochondrial stress tests were performed with the Seahorse XFe96/XF Pro FluxPak (Agilent, Cat# 103793-100) and 12 replicates per condition, according to the Seahorse XF Cell Mito Stress Test user manual. A375 cells were first plated on 96-well Fe96/XF Pro Cell Culture Microplates (Agilent, Cat#103794-100) at 5×10^3^ cells/well in complete DMEM (200 µL), incubated at 37°C (5% CO_2_, humidified atmosphere) overnight, then treated with or without JQ1 for 48 h. The XFe96/XF Pro sensor cartridge’s calibration was triggered 24 h before data acquisition by filling the calibrant plate with Seahorse XF Calibrant Solution and incubating at 37°C without CO_2_. On acquisition day, the medium was replaced with assay medium (Seahorse XF DMEM supplemented with 10 mM glucose, 1 mM pyruvate, 2 mM glutamine, and 1× Penicillin/Streptomycin, pH 7.5) and the plate was incubated for 2 h at 37°C without CO_2_. The XFe96/XF Pro sensor cartridge was then correctly filled with XF DMEM containing the following inhibitors: oligomycin (3.75 µM), carbonyl cyanide-4 (trifluoromethoxy) phenylhydrazone (2.75 µM), an antimycin A (3.0 µM)/rotenone (4.5 µM) mix, and 2-deoxy-D-glucose (50 mM; concentrations are those obtained after injection). Assays were run on a Seahorse XFe96 Analyzer (Agilent, Cat#S7800AR). The baseline oxygen consumption and the extracellular acidification rates were measured in basal conditions, followed by sequential injections of the inhibitors at the time points suggested by the manufacturer. Three measurements were performed after each inhibitor was injected. The results were normalized to the cell population, which was quantified immediately after the Seahorse run using the CyQUANT Cell Proliferation Assay (Invitrogen, Cat#C7026) according to the manufacturer's protocol. CyQUANT assay fluorescence intensities were detected on a Synergy H1 Hybrid Multi-Mode Reader (BioTeK, Cat# 11-120-531).

**Peptide arrays**

Residues corresponding to all previously reported phosphosites for BRD2, BRD3, and BRD4 in the phosphositeplus.org repository or those containing putative MK2 motifs (e.g., R-X-X-S) were synthesized as 13-mer peptides and arrayed on a membrane. For each site, the WT peptide sequences and corresponding non-phosphorylatable Ala-containing counterparts (either on the central Ser/Thr or on all other phosphorylatable residues) were synthesized on a MultiPep Automated Parallel Peptide Synthesizer using standard Fmoc chemistry and membrane handling as in (Dionne et al., 2018). Phosphorylation was assessed after incubating the membranes with human MK2 after TAP purification from K562 cells, in the presence of γ-^32^P-ATP as per (Warner et al., 2008). Radioactive ATP incorporation was detected by autoradiography on an Amersham Typhoon imager.

**QUANTIFICATION AND STATISTICAL ANALYSIS**

**CUT&RUN data analysis**

Sequencing results were first processed for data cleaning and quality assessment using Trim Galore (v0.6.4)/Cutadapt (v2.8) (Bioinformatics, 2021; Kechin et al., 2017), using standard parameters for paired-end sequencing, and FastQC (v0.11.9)/MultiQC (v1.11) (Ewels et al., 2016; S., 2010), using standard parameters. Cleaned sequences were aligned with the GRCh38 human reference genome using BowTie2 (v2.4.4) (Langmead and Salzberg, 2012) and standard parameters. Samtools (v1.15.1; (Li et al., 2009)) was used to sort samples by name and duplicates were marked using samblaster (v0.1.26) (Faust and Hall, 2014). Both unnormalized and bins per millions mapped reads (BPM)-normalized coverage was processed using bamCoverage (v3.3.0) (Ramirez et al., 2016). After sequence alignment, peaks were called against the corresponding IgG sample with Macs2 (v2.2.8; (Zhang and Su, 2012)) using the human genome size (-g hs) and a *q*-value of 0.05. Peaks were then imported into R and annotated using ChIPseeker (v1.38.0) (Yu et al., 2015), using limits of 3,000 bp upstream and downstream of a transcriptional start site for annotation using the Ensembl human genome database. The appropriate peak size (narrow or broad) was selected for each sample. A spiked-in *Saccharomyces cerevisiae* genome was aligned to the R64-1-1 reference genome using the same method to normalize the signal between samples.

**RNA-Seq**

Sequencing results were processed using a standard pipeline, including data cleaning and quality assessment using Trim Galore (v0.6.4)/Cutadapt (v2.8) (Bioinformatics, 2021; Kechin et al., 2017) using standard parameters for paired-end sequencing, and FastQC (v0.11.9)/MultiQC (v1.11) (Ewels et al., 2016; S., 2010), using standard parameters. Cleaned sequences were then aligned with the Gencode v27 human reference transcriptome using Kallisto (v0.46.1)(Bray et al., 2016) and the genomebam standard parameters. The results were then imported into R using tximport (v1.30.0) (Soneson et al., 2015) and differential expression was assessed using DESeq2 (v1.42.1) (Love et al., 2014). We used standard parameters and only included genes with at least three samples with > 5 normalized counts. Genes with fold change values > 1.5 and adjusted *p*-values < 0.05 were considered differentially expressed. Differentially expressed genes were subjected to gene ontology term enrichment analysis using the online Metascape portal between August 2023 and April 2024 (Zhou et al., 2019). Heatmaps were produced with ComplexHeatmap (v2.18.0) (Gu, 2022; Gu et al., 2016) using Ward.D2 clustering for both genes and samples and expression data transformed in log_2_(X+1). The gene set enrichment analysis (GSEA) tool (v.4.3.2)(Mootha et al., 2003) was used to examine enrichment in the “Hallmarks” gene set (h.all.v2023.1.Hs.symbols.gmt), using standard parameters and 1,000 permutations. Principal component analyses were performed with FactoMineR (v2.10)/FactoExtra (v1.0.7).

**CRISPR screen data analysis**

The CRISPR screen results were analyzed using the Model-based Analysis of Genome-wide CRISPR/Cas9 Knockout method as in Li *et al*. (Li et al., 2014).
