## Supplementary figures and images for "Systematic profiling of the acetyl lysine machinery reveals a role for MAPKAPK2 in bromodomain inhibitor resistance"

### Supplementary Figure S1

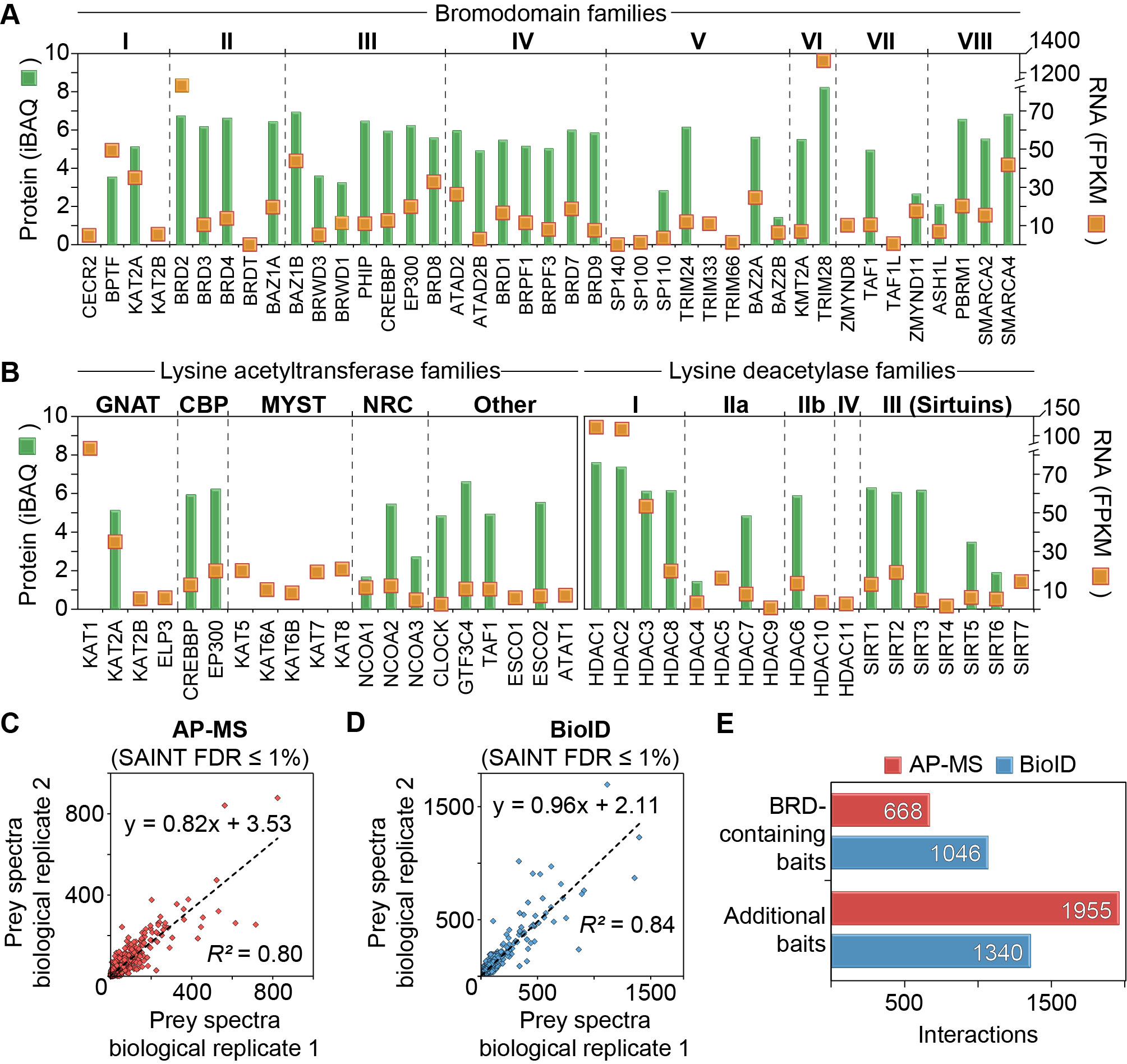

### Supplementary Figure S2

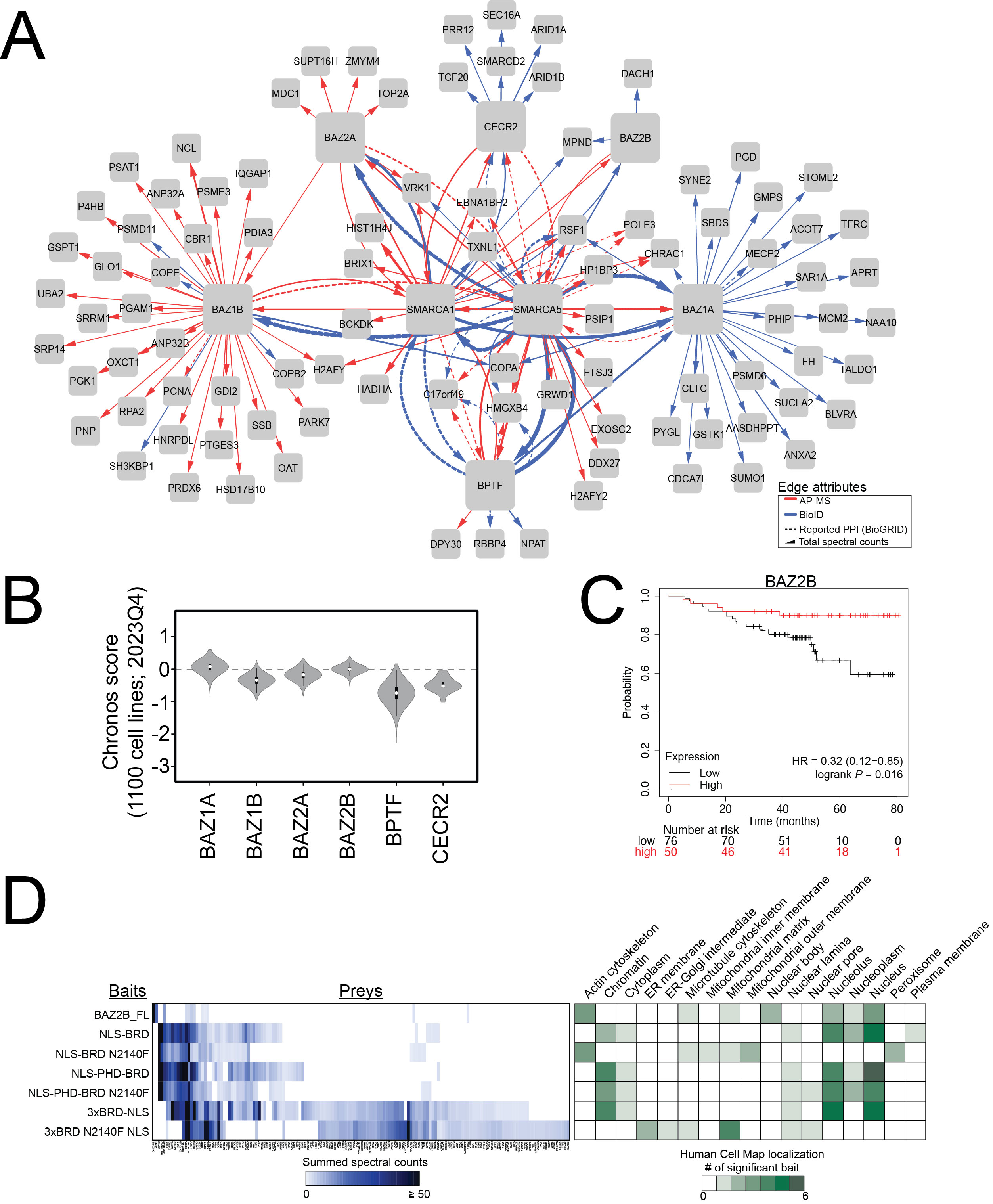

### Supplementary Figure S3

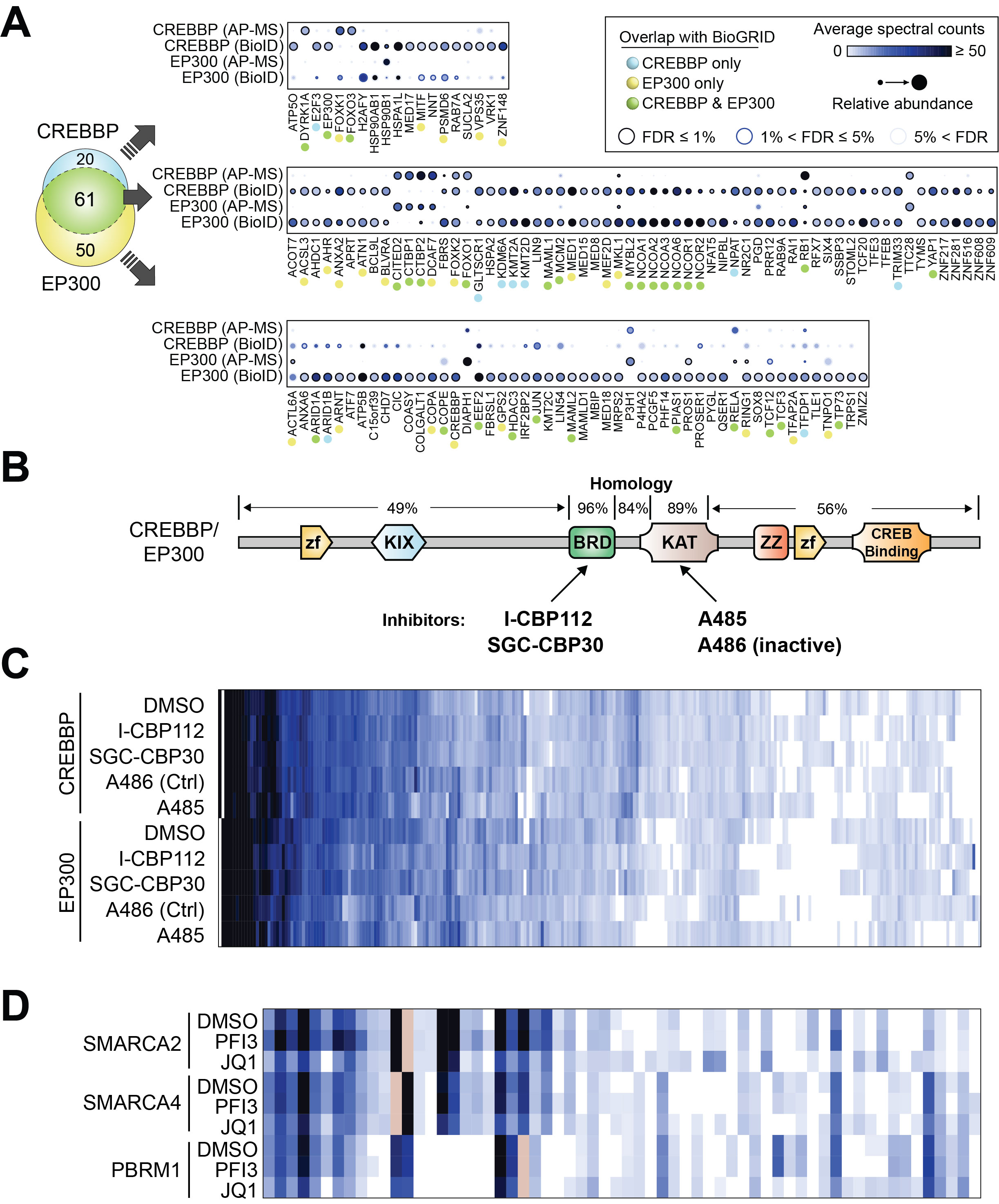

### Supplementary Figure S4

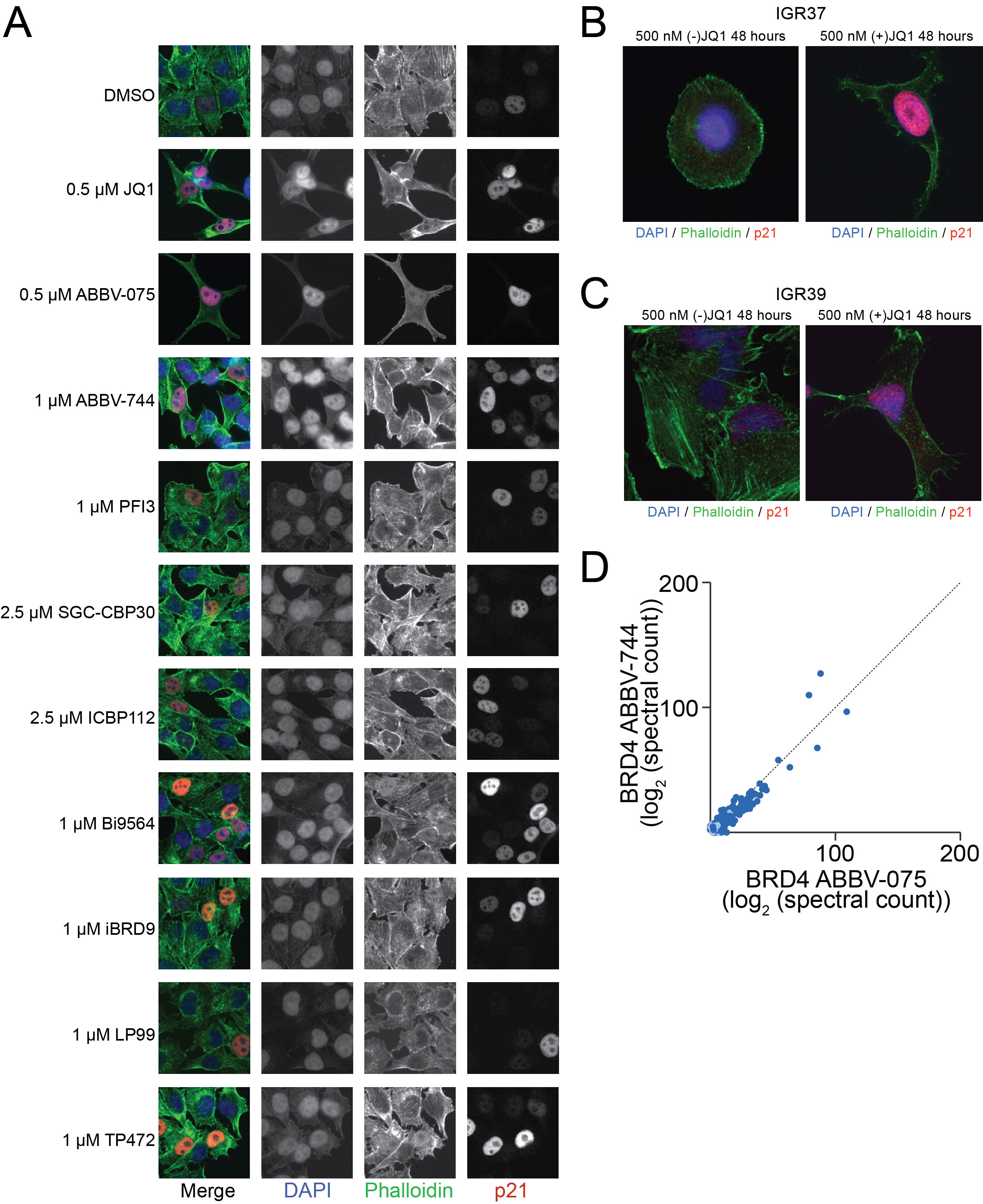

### Supplementary Figure S5

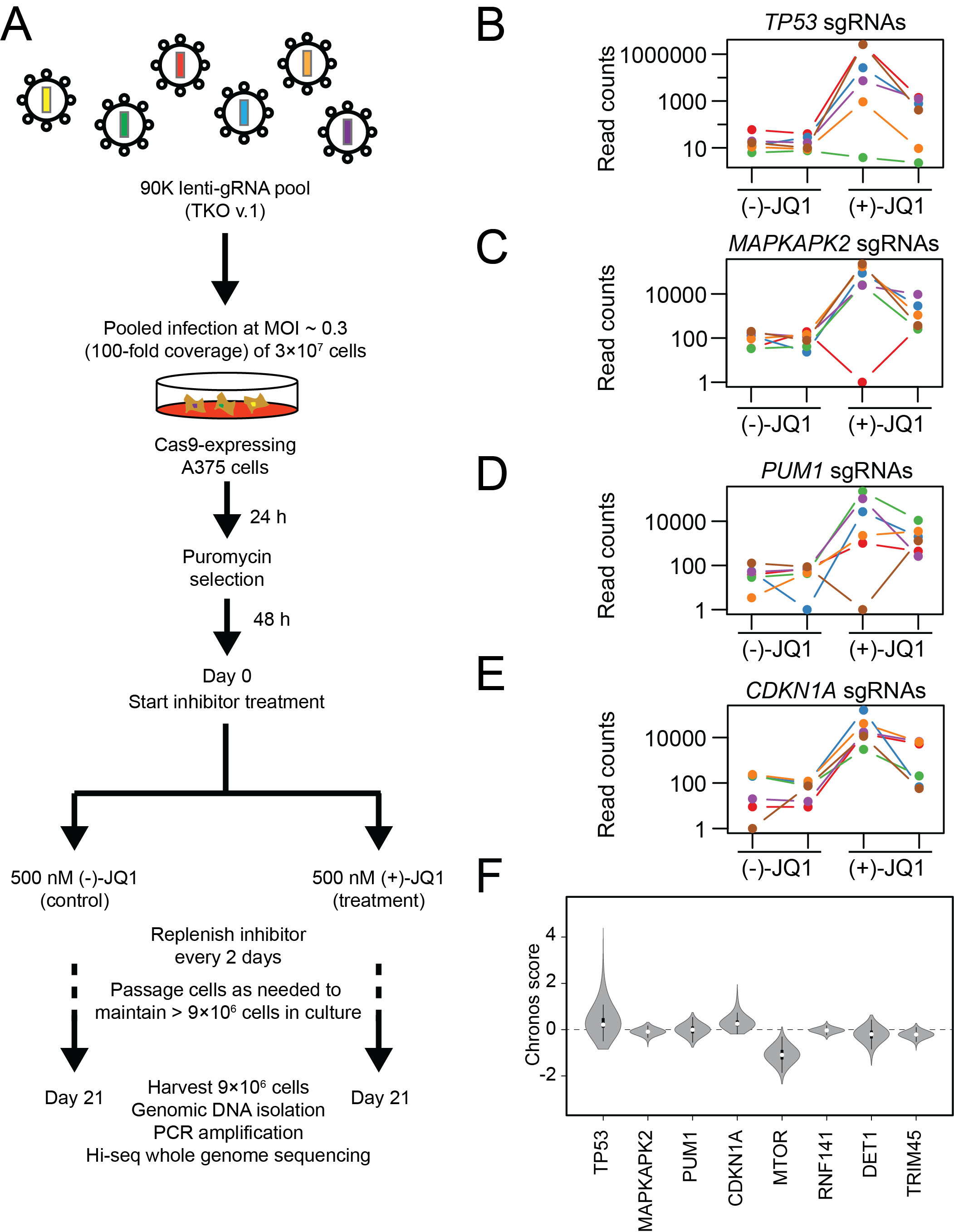

### Supplementary Figure S6

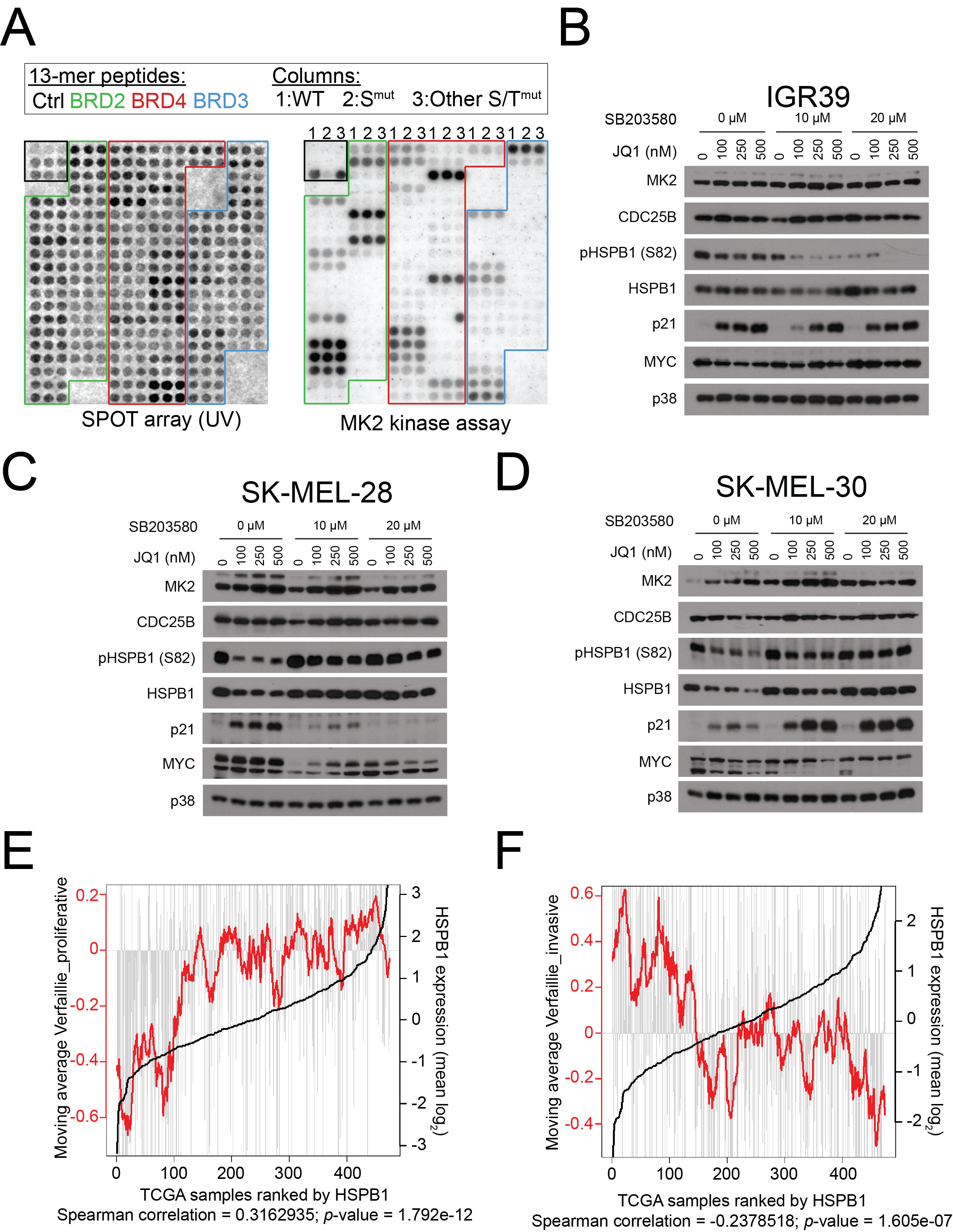

### Supplementary Figure S7

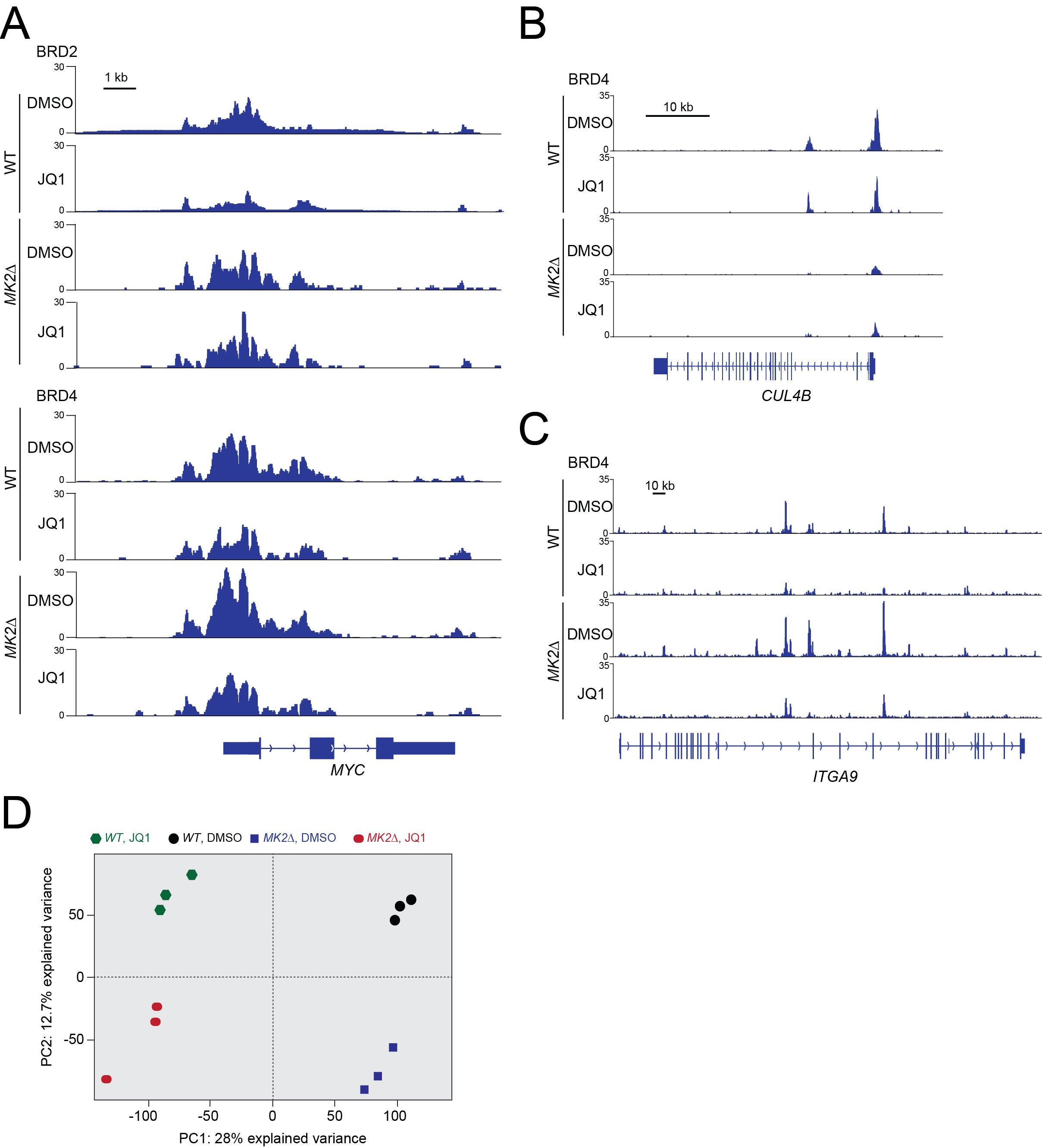
